# Ion-channel degeneracy and heterogeneities in the emergence of complex spike bursts in CA3 pyramidal neurons

**DOI:** 10.1101/2022.06.30.498226

**Authors:** Rituparna Roy, Rishikesh Narayanan

**Affiliations:** Cellular Neurophysiology Laboratory, Molecular Biophysics Unit, Indian Institute of Science, Bangalore, India

**Keywords:** complex spike bursts, hippocampus, computation, dendrite, ion channels, degeneracy

## Abstract

Complex spike bursting (CSB) is a characteristic electrophysiological signature exhibited by several neurons and has been implicated in neural plasticity, learning, perception, anesthesia, and active sensing. Here, we address the question of how pronounced intrinsic and synaptic heterogeneities affect CSB, with hippocampal CA3 pyramidal neurons (CA3PN) as a substrate where CSB emergence and heterogeneities are well-characterized. We randomly generated 12,000 unique models and found 236 valid models that satisfied 11 characteristic CA3PN measurements. These morphologically and biophysically realistic valid models accounted for gating kinetics and somato-dendritic expression profiles of 10 active ion channels. This heterogeneous population of valid models was endowed with broad distributions of underlying parameters showing weak pair-wise correlations. We found two functional subclasses of valid models, intrinsically bursting and regular spiking, with significant differences in the expression of calcium and calcium-activated potassium conductances. We triggered CSB in all 236 models through different intrinsic or synaptic protocols and observed considerable heterogeneity in CSB propensity and properties spanning models and protocols. Finally, we employed virtual knockout analyses and showed that synergistic interactions between intrinsic and synaptic mechanisms regulated CSB emergence and dynamics. Specifically, although there was a dominance of calcium and calcium-activated potassium channels in the emergence of CSB, individual deletion of none of the several ion channels or *N*-methyl-D-aspartate receptors resulted in the complete elimination of CSB across all models. Together, our analyses critically implicate ion-channel degeneracy in the robust emergence of CSB and other characteristic signatures of CA3PNs, despite pronounced heterogeneities in underlying intrinsic and synaptic properties.

## INTRODUCTION

Complex spike bursting (CSB) is a signature physiological response, elicited by several neuronal subtypes across different brain regions, typically in response to strong depolarizing inputs. Several protocols have been employed to trigger CSB, including coincident activation of different synaptic pathways (Wang *et al*., 2000; Takahashi & Magee, 2009; Xu *et al*., 2012; Bittner *et al*., 2015), synchronous synaptic inputs and opposing inhibition (Bastian & Nguyenkim, 2001; Harris *et al*., 2001; Royer *et al*., 2012; Grienberger *et al*., 2014; Raus Balind *et al*., 2019), coincidence of synaptic inputs and action potentials (Magee & Johnston, 1997; Larkum *et al*., 1999; Stuart & Hausser, 2001), and somatic and/or dendritic current injections (to elicit intrinsic CSB). A multitude of regulatory mechanisms have been implicated in the emergence of these CSB including dendritic size and arborization (Mainen & Sejnowski, 1996; Krichmar *et al*., 2002; Narayanan & Chattarji, 2010; van Elburg & van Ooyen, 2010), different combinations of intrinsic properties involved in intrinsic CSB (Wong & Prince, 1978; Hablitz & Johnston, 1981; Traub *et al*., 1991; Migliore *et al*., 1995; Golding *et al*., 1999; Williams & Stuart, 1999; Su *et al*., 2001; Lazarewicz *et al*., 2002; Wolfart & Roeper, 2002; Perez-Reyes, 2003; Swensen & Bean, 2003; Krahe & Gabbiani, 2004; Yue & Yaari, 2004; Swensen & Bean, 2005; Sipila *et al*., 2006; Vervaeke *et al*., 2006; Cueni *et al*., 2008; Hemond *et al*., 2008; Tazerart *et al*., 2008; Xu & Clancy, 2008; Narayanan & Chattarji, 2010; Raus Balind *et al*., 2019; Vickstrom *et al*., 2020), synaptic properties (Bastian & Nguyenkim, 2001; Harris *et al*., 2001; Royer *et al*., 2012; Grienberger *et al*., 2014; Raus Balind *et al*., 2019), and astrocytic activation (Kadala *et al*., 2015; Morquette *et al*., 2015; Ashhad & Narayanan, 2016; Condamine *et al*., 2018; Ashhad & Narayanan, 2019). Although bursts were traditionally studied primarily with reference to reliable information transmission, selectivity in transmitted information, and synaptic plasticity (Lisman, 1997; Izhikevich *et al*., 2003; Krahe & Gabbiani, 2004; Metzen *et al*., 2016; Sakmann, 2017), CSB and associated dendritic plateau potentials have received renewed attention with recent demonstrations that have implicated them in behavioral time scale synaptic plasticity (Bittner *et al*., 2015; Bittner *et al*., 2017; Magee & Grienberger, 2020; Zhao *et al*., 2020), perception (Larkum, 2013; Manita *et al*., 2015; Takahashi *et al*., 2016; Takahashi *et al*., 2020), anesthesia (Aru *et al*., 2020; Redinbaugh *et al*., 2020; Suzuki & Larkum, 2020), active sensing (Lavzin *et al*., 2012; Xu *et al*., 2012; Ranganathan *et al*., 2018), and learning (Doron *et al*., 2020; Larkum *et al*., 2022). Despite these broad physiological implications, the question of how neurons generate CSB despite widespread heterogeneities in their synaptic and intrinsic properties has remained unexplored.

Cornu Ammonis 3 (CA3) is a subregion of the hippocampus with characteristic recurrent connectivity among pyramidal neurons that manifest signature burst firing (Wong & Prince, 1978; Hablitz & Johnston, 1981; Traub, 1982; Bilkey & Schwartzkroin, 1990; Traub *et al*., 1991; Migliore *et al*., 1995; Migliore *et al*., 1999; Lazarewicz *et al*., 2002; Hemond *et al*., 2008; Narayanan & Chattarji, 2010; Zeldenrust *et al*., 2018). The CA3 receives inputs from the dentate gyrus and the entorhinal cortex, and has been implicated in spatial navigation, spatial memory, pattern completion, and episodic memory (Nakazawa *et al*., 2002; Andersen *et al*., 2006; Kesner, 2007; Miles *et al*., 2014). CA3 pyramidal neurons elicit CSB when stimulated with a multitude of stimulus paradigms employ strong depolarizing inputs to the neuron as a common substrate (Raus Balind *et al*., 2019). CSB is characterized by several consecutive action potentials (AP) within a very small time-interval riding on top of a voltage ramp, with AP amplitude reducing within the burst, typically followed by a prolonged period of slow after-hyperpolarization.

In this study, we explore several signature electrophysiological characteristics of CA3 pyramidal neurons, including CSB elicited through multiple protocols, and ask if this constellation of characteristic electrophysiological measurements constrain the biophysical composition of CA3 pyramidal neurons. A single hand-tuned model wouldn’t be sufficient to accommodate physiologically observed parametric and measurement heterogeneities or to effectively address the question on the role of neural heterogeneities on CSB emergence. Therefore, we employed an unbiased stochastic search approach that has been effectively used in other neural systems (Foster *et al*., 1993; Prinz *et al*., 2004; Taylor *et al*., 2009; Rathour & Narayanan, 2012, 2014; Rathour *et al*., 2016; Migliore *et al*., 2018; Mittal & Narayanan, 2018; Mishra & Narayanan, 2019; Goaillard & Marder, 2021; Mishra & Narayanan, 2021a) to explore the parametric space required for effectively achieving the characteristic properties of CA3 pyramidal neurons. The overall goal here was to understand the biophysical basis of CSB emergence in CA3 pyramidal neurons, while carefully accounting for the widespread heterogeneities in the parametric and measurement spaces.

Our extensive search process involving 12,000 morphologically realistic conductance-based models yielded 236 models that manifested signature characteristics of CA3 pyramidal neurons, which were either regular spiking or intrinsically bursting neurons. Analyses of these two classes of models unveiled a dominant role for calcium and calcium-activated potassium channels in mediating the distinction between these two classes. We subjected these models to five different protocols for eliciting CSB and studied the different signature measurements from these recordings. Our analyses of model measurements and parameters unveiled the expression of ion-channel degeneracy in CA3 pyramidal neurons in the manifestation of characteristic physiological properties including CSB.

## METHODS

### CA3b pyramidal neuron model

A morphologically realistic neuron model (Fig. 1*A*) was constructed from the three-dimensional reconstruction of a CA3b pyramidal neuron (Cannon *et al*., 1998) with 10 different types of ion channels whose gating kinetics were adopted from previous literature (Migliore *et al*., 1995). The passive parameters were maintained constant across the entire neuron with the specific membrane capacitance set at 0.75 μF/cm^2^, specific membrane resistivity *R*_m_ was 60 kΩ.cm^2^, and axial resisitivity *R*_a_ was assigned as 200 Ω.cm (default values in the base model are provided). The neuronal model was compartmentalized according to the *d*_λ_ rule (Carnevale and Hines, 2006) such that iso-potential condition was maintained within each compartment. The ion channels that were incorporated into the model were a fast sodium (NaF) channel; three voltage-gated potassium channels: delayed rectifier (KDR), *A* type (KA), and *M* type (KM); three voltage-gated calcium channels: *L* type (CaL), *N* type (CaN) and *T* type (CaT); two calcium-dependent potassium channels: small-conductance (SK) and big-conductance (BK); and hyperpolarization activated cyclic nucleotide-gated (HCN) non-specific cation channel.

**Figure 1:**
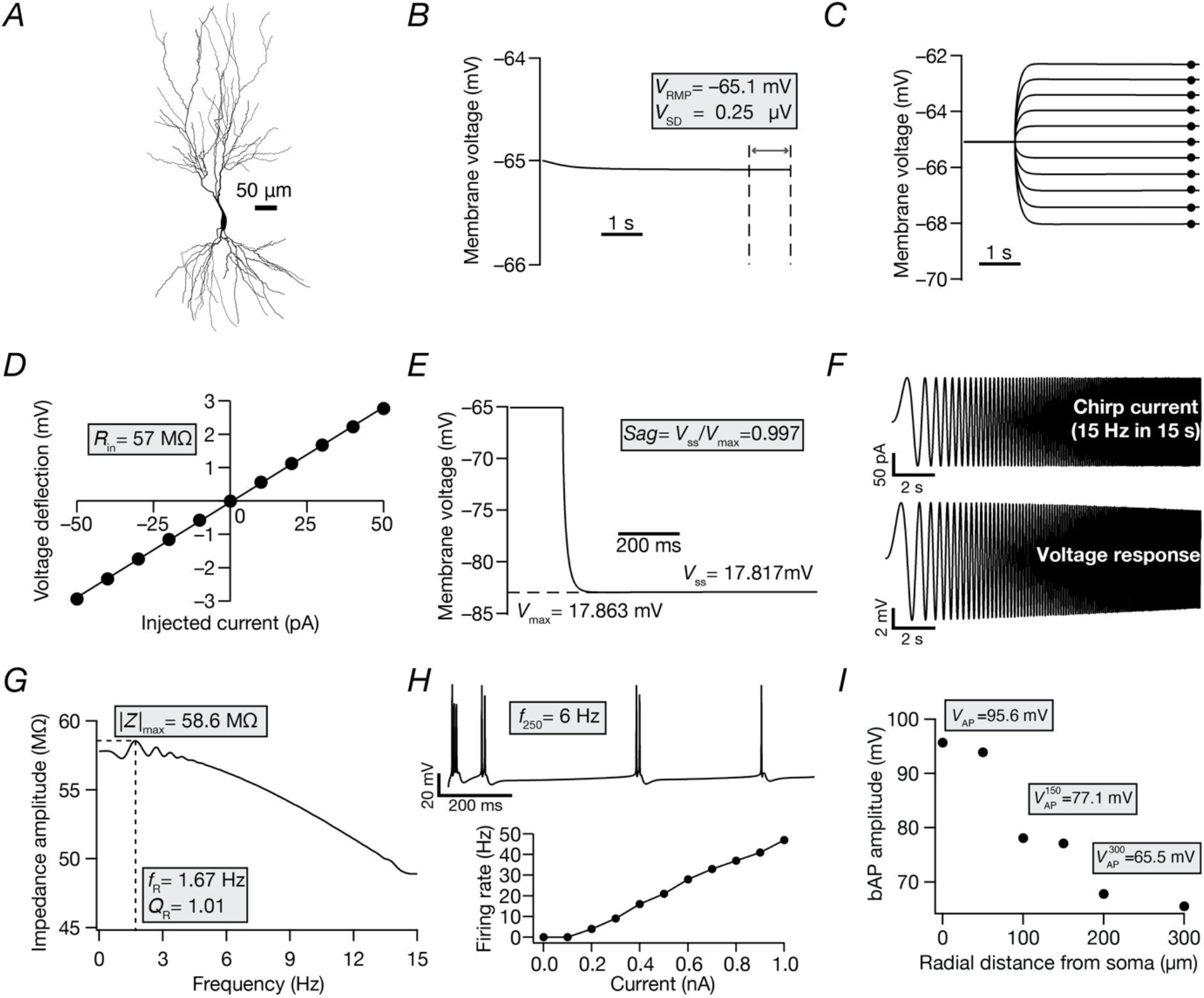
The base model satisfied characteristic intrinsic somato-dendritic electrophysiological measurements of CA3 pyramidal neurons. *A*, 2D projection of a reconstructed 3D hippocampal pyramidal CA3b neuron model used as the base model. *B–I*, The 11 electrophysiological measurements used to characterize the morphologically realistic model of CA3b pyramidal neuron, highlighted in gray boxes. *B*, Resting membrane potential (*V_RMP_*) was calculated as the mean value of recorded voltage, in the absence of injected current, within the 5 s and 6 s window. The standard deviation (*V_SD_*) was computed from the same trace spanning the 5–6 s period. All subsequent measurements were performed after an initial delay of 5 s, to allow *V_RMP_* to reach steady-state. *C*, Input resistance (*R_in_*) was calculated from the voltage responses elicited by injecting current pulses of amplitude –50 pA to +50 pA in steps of 10 pA for 300 ms. *D*, *V–I* plot showing steady-state voltage responses *vs*. injected current values, obtained from traces in panel *C*. *R_in_* was computed as the slope of the linear fit to this *V-I* plot. *E*, Sag ratio (*Sag*) was measured as the ratio of the steady-state membrane potential deflection (*V_SS_*) to the peak membrane potential deflection (*V_max_*) in the voltage response to a hyperpolarizing current pulse of 250 pA injected for a duration of 800 ms. *F*, Impedance based measurements were obtained from the voltage response (*Bottom*) to a chirp current stimulus (*Top*). *G*, Impedance amplitude profile depicting the resonance frequency (*f_R_*) at which the maximal response (|*Z*|_*max*_) was elicited by the neuron. Resonance strength (*Q_R_*) was calculated as the ratio of impedance amplitude at *f_R_* to the impedance amplitude at 0.5 Hz. *H*, *Top*, voltage response to a current pulse of 250 pA injected for 1 s. The firing rate (*f*_250_) was calculated as the number of action potentials (AP) elicited during the period of current injection. *Bottom*, AP firing rate plotted as a function of injected pulse-current (1 s duration) amplitude. *I*, Backpropagating action potential (bAP) amplitude plotted as a function of radial distance from the soma. Recordings were obtained at different dendritic locations along the somato-dendritic axis after somatic injection of a depolarizing current of 1 nA for 50 ms. The amplitude of the propagating AP at various locations were measured from the respective recordings. 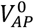: AP amplitude at soma. 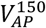: bAP amplitude at a dendrite around 150 μm from the soma. 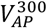: bAP amplitude at a dendrite around 300 μm from the soma.

All ion channels were modeled using Hodgkin-Huxley dynamics (Hodgkin & Huxley, 1952). Sodium and potassium channel currents were modeled using an Ohmic formulation with reversal potentials set at *E*_Na_ = 50 mV and *E*_K_ = −91 mV. Current through calcium channels was modeled using the GHK formulation (Goldman, 1943; Hodgkin & Katz, 1949; Migliore *et al*., 1995). A mobile calcium buffer mechanism and a calcium pump mechanism, both uniformly distributed across the somato-dendritic arbor, accounted for calcium decay (Migliore *et al*., 1995). The total calcium buffer concentration and the total pump density were model parameters that governed calcium dynamics in the model. Longitudinal and radial diffusion (across 4 annuli) of calcium, the unbound and bound buffers were incorporated into the model definition.

The distribution for all ion channels followed available computational and experimental data from CA3 pyramidal neurons (Migliore *et al*., 1995; Narayanan & Chattarji, 2010; Kim *et al*., 2012). NaF, KDR, CaN, SK, BK, and CaT channels were uniformly distributed in the soma and along the dendritic arbor (Migliore *et al*., 1995; Lazarewicz *et al*., 2002; Narayanan & Chattarji, 2010). The distribution of CaL channels was perisomatic (≤ 50 μm from the soma) (Migliore *et al*., 1995; Narayanan & Chattarji, 2010). Different kinetic schemes were used for the proximal (≤100 μm from the soma) and distal distribution (>100 μm) of the KA channel (Migliore *et al*., 1999; Narayanan & Chattarji, 2010). Our models did not have an explicit axonal compartment (Narayanan & Chattarji, 2010). *M*-type potassium channel was distributed peri-somatically (≤ 100 μm) and the kinetics was adopted from earlier studies (Migliore *et al*., 1995; Lazarewicz *et al*., 2002). All somatodendritic active and passive parameters were tuned to match the distance dependent electrophysiological measurements for backpropagating action potential, bAP (Kim *et al*., 2012) for the CA3b pyramidal neuron. Although there are lines of evidence for the density of HCN channels in CA3 pyramidal neurons to be lower compared to their CA1 counterparts, the sub-cellular distribution of HCN along the CA3 dendritic arbor is unclear (Raus Balind *et al*., 2019). Therefore, the general distribution pattern following that of CA1 pyramidal neurons was adopted, with increasing density along the apical dendrite. HCN channel was distributed in a way that its density increased as a function of radial distance *x* from the soma along the apical dendrite (Lazarewicz *et al*., 2002), and followed the kinetics as mentioned (Magee, 1998).

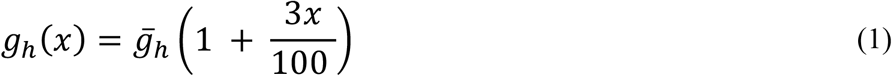

All model parameters and their respective base values are listed in Table 1.

**Table 1:** Model parameters with their base values and the range spanned for the unbiased stochastic search

|  | Parameter(unit) | Symbol | Base value | Range |
| --- | --- | --- | --- | --- |
| **Passive properties (uniform across the neuron)** |  |  |  |  |
| 1 | Membrane resistance ($\text{k}\Omega\cdot\text{cm}^{-2}$) | $R_m$ | 60 | 60–100 |
| 2 | Axial resistance ($\Omega\cdot\text{cm}$) | $R_a$ | 200 | 150–400 |
| **Active properties** |  |  |  |  |
| **Spike-generating channels (uniformly distributed)** |  |  |  |  |
| 3 | Maximal conductance for NaF ($\text{mS}\cdot\text{cm}^{-2}$) | $\bar{g}_{Na}$ | 15 | 7.5–30 |
| 4 | Maximal conductance for KDR ($\text{mS}\cdot\text{cm}^{-2}$) | $\bar{g}_{KDR}$ | 11 | 5.5–22 |
| **HCN channel (increase from soma to distal apical dendrite)** |  |  |  |  |
| 5 | Maximal conductance ($\mu\text{S}\cdot\text{cm}^{-2}$) | $\bar{g}_h$ | 1 | 0.5–6 |
| **A-type potassium channel (linear increase with distance along apical dendrite)** |  |  |  |  |
| 6 | Maximal conductance ($\text{mS}\cdot\text{cm}^{-2}$) | $\bar{g}_{KA}$ | 0.1 | 0.05–0.2 |
| **L-type calcium channel (peri-somatic distribution)** |  |  |  |  |
| 7 | Maximal conductance ($\text{mS}\cdot\text{cm}^{-2}$) | $\bar{g}_{CaL}$ | 2.5 | 1.25–5 |
| **T-type calcium channel (uniformly distributed)** |  |  |  |  |
| 8 | Maximal conductance ($\text{mS}\cdot\text{cm}^{-2}$) | $\bar{g}_{CaT}$ | 0.25 | 0.125–0.5 |
| **N-type calcium channel (uniformly distributed)** |  |  |  |  |
| 9 | Maximal conductance ($\text{mS}\cdot\text{cm}^{-2}$) | $\bar{g}_{CaN}$ | 2.5 | 1.25–5 |
| **M-type potassium channel (peri-somatic distribution)** |  |  |  |  |
| 10 | Maximal conductance ($\text{mS}\cdot\text{cm}^{-2}$) | $\bar{g}_{KM}$ | 0.01 | 0.005–0.02 |
| **Large conductance calcium-activated potassium BK channel (uniformly distributed)** |  |  |  |  |
| 11 | Maximal conductance ($\text{mS}\cdot\text{cm}^{-2}$) | $\bar{g}_{BK}$ | 0.8 | 0.4–1.6 |
| **Small conductance calcium-activated potassium SK channel (uniformly distributed)** |  |  |  |  |
| 12 | Maximal conductance ($\text{mS}\cdot\text{cm}^{-2}$) | $\bar{g}_{SK}$ | 0.5 | 0.25–1 |
| **Calcium handling mechanisms** |  |  |  |  |
| 13 | Total calcium pump | totpp | 0.2 | 0.01–0.5 |
| 14 | Total calcium buffer | totbuf | 1.2 | 0.1–2.5 |

### Intrinsic measurements

Models were validated against an array of 11 characteristic electrophysiological measurements (sub- and supra-threshold) from CA3 pyramidal neurons (Figure 1, Table 2). These measurements were computed using well-established procedures (Narayanan & Johnston, 2007, 2008; Rathour & Narayanan, 2014; Basak & Narayanan, 2018; Mittal & Narayanan, 2018; Mishra & Narayanan, 2020; Seenivasan & Narayanan, 2020) and are described below. Resting membrane potential (*V*_RMP_) was calculated as the mean value of recorded voltage within the 5–6 s window from the 6-s recordings performed in the absence of injected current (Fig. 1*B*). The standard deviation (*V_SD_*) was computed from the same trace spanning the 5–6 s period. All sub- and supra-threshold measurements were performed after an initial delay of 5 s, to allow *V*_RMP_ to reach steady-state value. Input resistance (*R_in_*) was calculated from the voltage responses elicited by injecting current pulses of amplitude −50 pA to +50 pA in steps of 10 pA for 300 ms (Fig. 1*C*). *R_in_* was computed as the slope of the linear fit to a *V–I* plot of the steady-state voltage responses *vs*. injected current values (Fig. 1*D*). Sag ratio (*Sag*) was measured as the ratio of the steady-state membrane potential deflection (*V_SS_*) to the peak membrane potential deflection (*V_max_*) in the voltage response to a hyperpolarizing current pulse of 250 pA injected for a duration of 800 ms (Fig. 1*E*).

**Table 2:** Intrinsic measurements of CA3 pyramidal neurons and their respective electrophysiological bounds. These measurements were derived from electrophysiological recordings reported in (Hemond *et al*., 2008; Kim *et al*., 2012; Raus Balind *et al*., 2019).

|  | Measurement | Symbol | Lower bound | Upper bound |
| --- | --- | --- | --- | --- |
| 1 | Resting membrane potential (mV) | $V_{RMP}$ | -70 | -60 |
| 2 | Standard deviation (μV) | $V_{SD}$ | 0 | 0.01 |
| 3 | Input resistance ($M\Omega$) | $R_{in}$ | 60 | 150 |
| 4 | Sag ratio | $Sag$ | 0.94 | 1 |
| 5 | Resonant frequency (Hz) | $f_R$ | 1 | 1.76 |
| 6 | Strength of resonance | $Q_R$ | 1 | 1.1 |
| 7 | Impedance amplitude ($M\Omega$) | $\vert Z\vert _{max}$ | 65 | 150 |
| 8 | Firing rate at 250 pA (Hz) | $f_{250}$ | 4 | 10 |
| 9 | AP amplitude at soma (mV) | $V_{AP}^0$ | 90 | 110 |
| 10 | AP amplitude at 150 μm (mV) | $V_{AP}^{150}$ | 70 | 90 |
| 11 | AP amplitude at 300 μm (mV) | $V_{AP}^{300}$ | 60 | 80 |

Impedance-based measurements were obtained from the voltage response to a chirp current stimulus (Fig. 1*F*). The chirp stimulus employed here was a constant-amplitude (100 pA peak-to-peak) sinusoidal current with frequency increasing linearly from 0–15 Hz over a period of 15 s (Narayanan & Johnston, 2007, 2008; Hemond *et al*., 2009; Basak & Narayanan, 2018). The Fourier transform of the voltage response was divided by the Fourier transform of the chirp current to obtain complex impedance *Z*(*f*), as a function of frequency *f*. Impedance amplitude profile |*Z*(*f*)| was computed as:

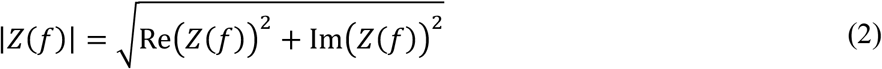

where Re(*Z*(*f*)) and *Im*(*Z*(*f*)) are the real and imaginary parts of the impedance *Z*(*f*), respectively. The frequency at which the value of |*Z*(*f*)| reached its maximum (|*Z*|_*max*_) was defined as resonance frequency *f_R_* (Fig. 1*G*). Resonance strength (*Q_R_*) was calculated as the ratio of impedance amplitude at *f_R_* to the impedance amplitude at 0.5 Hz.

The firing rate in response to a pulse current injection of 250 pA (*f*_250_) was calculated as the number of action potentials (AP) elicited during the 1-s period of current injection (Fig. 1*H*). To compute backpropagating action potential (bAP) amplitude as a function of distance from the soma, recordings were obtained at different dendritic locations along the somato-dendritic axis after somatic injection of a depolarizing current of 1 nA for 50 ms. The amplitude of the propagating AP at various locations was measured from the first AP in the respective recordings. 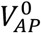: AP amplitude at soma. 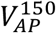: bAP amplitude at a dendrite ~150 μm from the soma. 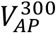: bAP amplitude at a dendrite ~300 μm from the soma (Fig. 1*I*).

### Multi-parametric multi-objective stochastic search (MPMOSS)

We performed MPMOSS on a 14-parameter space that spanned passive properties, ion channel conductance values, and calcium-handling mechanisms (Table 1). A total of 12,000 unique models were randomly generated by sampling each of these parameters from a uniform distribution that spanned the respective bounds (Table 1). Sub- and supra-threshold physiological measurements of each model were computed and were validated against their respective bounds obtained from electrophysiological recordings from CA3 pyramidal neurons (Table 2). Models with all 11 intrinsic measurements falling within their respective physiological bounds were declared valid (Fig. 2*A*). The parameters and the measurements of these valid models were then subjected to further analyses exploring heterogeneities and degeneracy.

**Figure 2:**
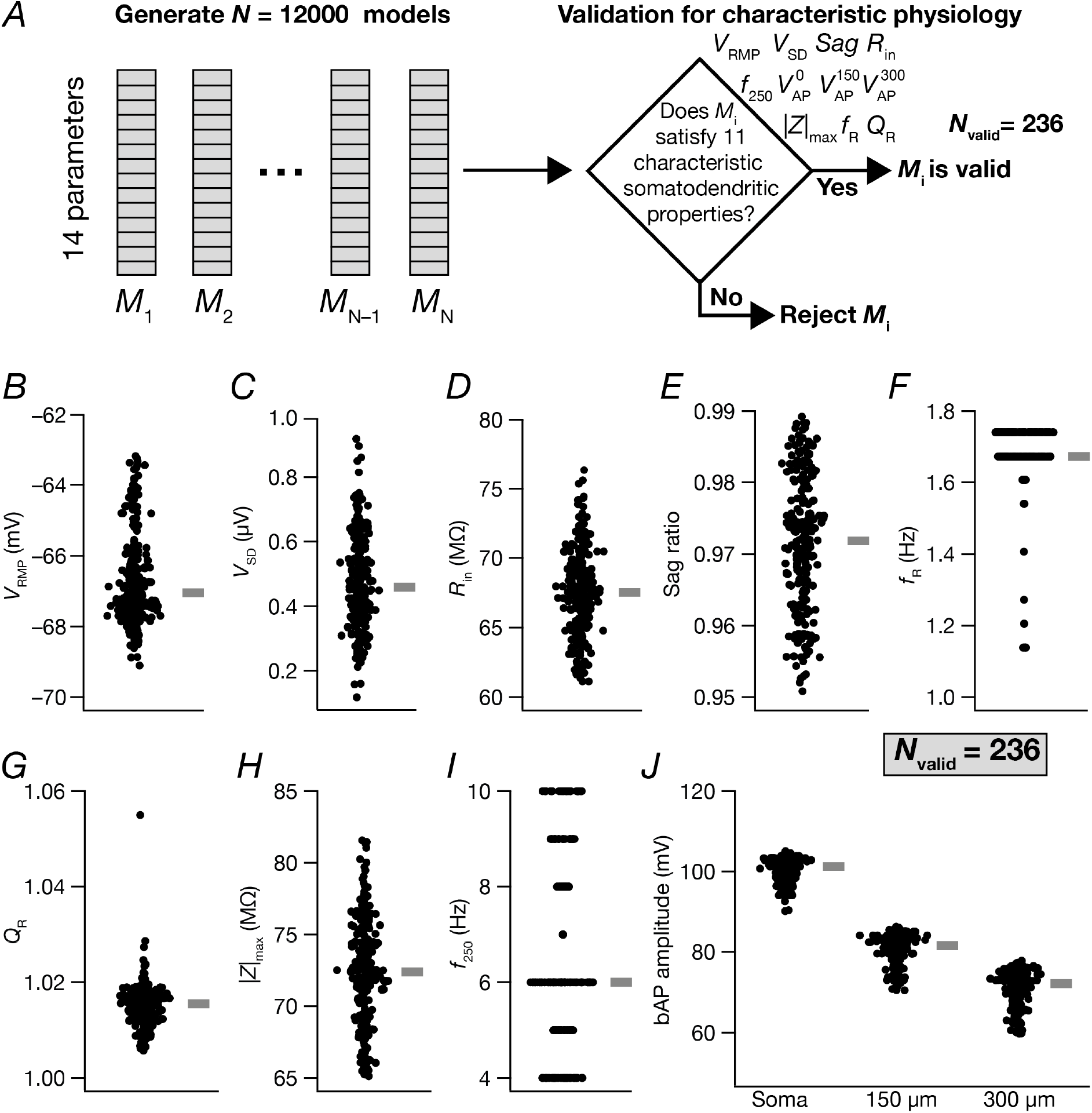
Heterogeneous distribution of signature somato-dendritic intrinsic measurements in physiologically valid CA3 pyramidal neuron models obtained from a multi-parametric multi-objective stochastic search. *A*, Illustration of the multi-parametric multi-objective stochastic search (MPMOSS) spanning 14 search parameters (Table 1) and 11 validation measurements (Table 2). *B–J*, Beeswarm plots depicting the heterogeneous distribution of the 11 intrinsic somato-dendritic measurements from each of the 236 physiologically valid models. All 11 measurements from each of the 236 valid models satisfied all physiological bounds characteristic of CA3 pyramidal neurons (Table 2). Thick lines on the right depict respective median values.

### Intrinsic bursting and regular spiking models

We delineated intrinsically bursting (IB) neurons from their regular spiking (RS) counterparts by analyzing the somatic voltage response to a current pulse of 240 pA injected into the soma for 5.5 s after an initial delay of 5 s (to allow RMP to settle to a steady-state value). Valid neurons derived from the MPMOSS procedure were placed in the IB class if there were ≥ 3 APs within 25 ms and if the action potential amplitude reduced with successive action potentials (quantified by the difference between the first and the second action potentials within the burst). These quantifications were arrived by visualizing the inter-spike interval (ISI) histogram of IB *vs*. RS neurons (Holt *et al*., 1996; Frerking *et al*., 2005; Selinger *et al*., 2007).

We performed both linear (principal component analysis, PCA) and non-linear (*t*-distributed stochastic neighbor embedding, *t*-SNE) dimensionality reduction techniques on the 12-dimensional active parametric space (Table 1) to understand the distributions of the underlying parameters governing the valid model population. These analyses were performed to assess differences in parametric distributions between the RS and the IB classes of neurons.

### Synaptic model

Co-localized AMPAR-NMDAR synapses were placed randomly in the *stratum radiatum* region along the apical dendrite of the CA3 pyramidal neuron. The number of synapses was set to a default value of 100. The default ratio of NMDAR:AMPAR was set to 2 and the ionic currents through these receptors were modeled using the GHK convention (Goldman, 1943; Hodgkin & Katz, 1949; Narayanan & Johnston, 2010). The current through NMDA receptors reflected their voltage-dependence and ionic composition, carried by sodium, potassium, and calcium ions:

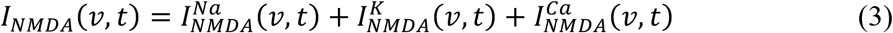

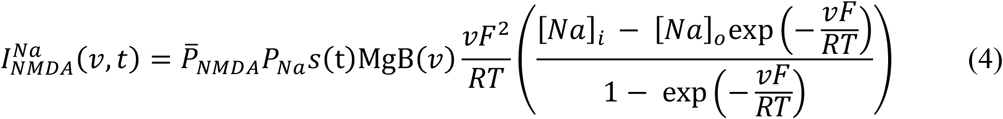

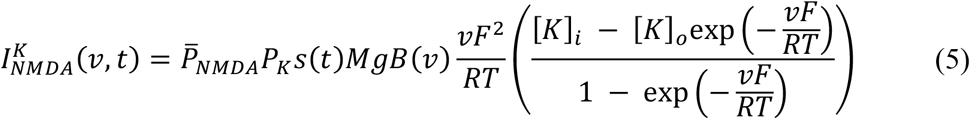

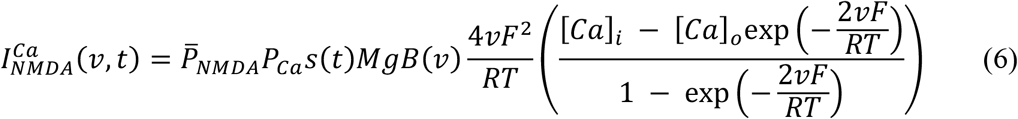

where 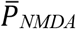 defined the maximum permeability of the NMDA receptor. The relative permeability values for Na^+^ and K^+^ were set as *P_Na_* = *P_K_* = 1 and for calcium, *P_Ca_* = 10.6. The intra- and extra-cellular concentrations for the different ions were set as: [*Na*]_*i*_ = 18 mM, [*Na*]_*o*_ = 140 mM, [*K*]_*i*_ = 140 mM, [*K*]_*o*_ = 5 mM, [*Ca*]_*i*_ = 100 nM and [*Ca*]_*o*_ = 2 mM. These ionic concentrations ensured that the reversal potentials for both AMPA and NMDA were set at 0 mV. The role of magnesium in regulating the activity of the NMDAR was accounted by the *MgB*(*v*) factor (Jahr & Stevens, 1990):

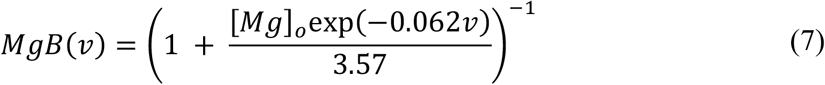

where [*Mg*]_*o*_ = 2 mM. *s*(*t*) in equations (4–6) governed the kinetics of the NMDA receptor as follows (Jahr & Stevens, 1990):

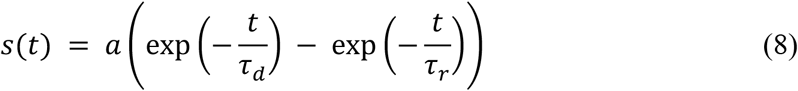

where *a* defined a normalization factor that ensured that 0 ≤ *s*(*t*) < 1. *τ_r_* (= 5 ms) and *τ_d_* (= 50 ms) represented the rise time and decay time constant, respectively.

The AMPAR current was modelled following the GHK convention, and was driven by two ions (sodium and potassium):

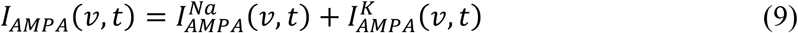

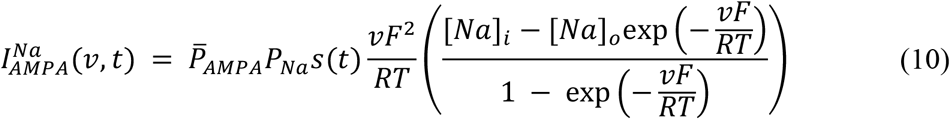

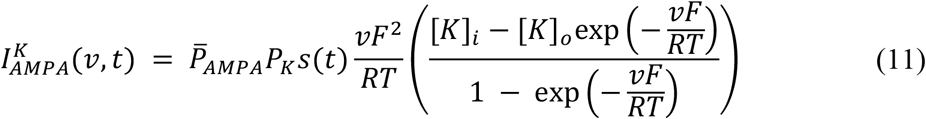

where 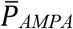 defined the maximum permeability of the AMPA receptor, with *P_Na_* = *P_K_* = 1. The rise and decay time constants of AMPAR were as *τ_r_* = 2ms and *τ_d_* = 10 ms. When present, these synapses were stimulated by a pre-synaptic spike train at 5 Hz frequency.

### Complex spike burst generation

We tried three different classes of protocols to study the emergence of complex spike bursts, adapting the strategies in electrophysiological studies (Raus Balind *et al*., 2019):

1. *Large somatic current injections:* Five pulses of large depolarizing currents with amplitude of 600 pA, 900 pA, or 1200 pA were injected into the cell body for a duration of 100 ms with an inter-pulse duration of 80 ms after the usual delay period of 5 s for the RMP to attain steady-state values.
2. *Large dendritic current injections:* Five pulses of large depolarizing currents with an amplitude of 1000 pA were injected into an apical dendritic location (~150 μm away from the soma) for a duration of 50 ms with an inter-pulse duration of 50 ms, after the usual delay period of 5 s for the RMP to attain steady-state values.
3. *Synaptic stimulation:* Synaptic inputs were given by synchronously stimulating 100 colocalized AMPAR-NMDAR synapses with pre-synaptic spike trains arriving at 5 Hz on dendritic locations randomly distributed over the *stratum radiatum*.

Complex spike bursts were identified across the 5 pulses in all the models using characteristic physiological measurements (Table 3): (1) ≥ 3 AP in 25 ms within the pulse; (2) Maximal reduction in the amplitude of the second AP relative to the first, assessed as the difference between the first AP and the second AP within each pulse; (3) ramp-like depolarization of the voltage across each pulse, calculated by subjecting the traces to a median filter of 0.5 s window to remove spikes. *V*_ramp_ was calculated as the difference of the peak value of this filtered trace from the resting membrane potential (Harvey *et al*., 2009; Basak & Narayanan, 2018; Roy & Narayanan, 2021). CSB rate was obtained in the range of 0–1 for all the different input configurations, as the ratio between the number of pulses that satisfied each of the CSB criteria and the total number of pulses (Raus Balind *et al*., 2019). Since the first pulse always demonstrated complex spiking burst for all the cases, CSB rate was computed by taking the remaining 4 pulses.

**Table 3:** Measurements and their bounds used to validate complex spike bursting.

|  | Measurement | Lower bound | Upper bound |
| --- | --- | --- | --- |
| 1 | Number of action potentials in the CSB | 3 | — |
| 2 | $\Delta V_{AP} = V_{AP}^1 - V_{AP}^2$ (mV) | 10 | 30 |
| 3 | Ramp amplitude, $V_{ramp}$ (mV) | 10 | 30 |

### Virtual knockout models

We used the virtual knockout models (VKM) paradigm (Rathour & Narayanan, 2014; Anirudhan & Narayanan, 2015; Mukunda & Narayanan, 2017; Basak & Narayanan, 2018; Mittal & Narayanan, 2018; Seenivasan & Narayanan, 2020; Mishra & Narayanan, 2021b; Roy & Narayanan, 2021) to identify the impact of individual ion channels involved in eliciting distinctive complex spike bursts in CA3 pyramidal neurons. For each of the 8 ion channel subtypes, we set the maximal conductance value to zero across all valid models and assessed the impact of such virtual knockout on different CSB measurements. We used the 900-pA somatic current injection and the synaptic stimulation protocols for inducing CSB and assessed voltage traces for all valid models, independently for each of the 8 individual virtual knockouts. We compared these outcomes with the CSB measurements from the respective base models to evaluate the impact of individual ion channels on each CSB measurement.

We assessed the impact of NMDAR virtual knockout on CSB measurements obtained with synaptic stimulation across all valid models by setting the NMDAR maximal permeability to zero during synaptic stimulation (equations 4–6). The AMPAR permeability was intact across models, thus allowing synaptic transmission to occur when the synaptic stimulation protocol for eliciting CSB was employed. To understand the role of calcium mediated through the NMDA receptors in triggering CSB in the neurons, we measured local calcium concentrations across pulses and plotted them in the presence and absence (virtual knockout) of NMDARs.

### Computational details

All simulations were performed using the NEURON programming environment (Carnevale and Hines, 2006) at 34° C. The simulation step size was set as 25 μs. Data analysis and plotting of graphs were done using custom-written codes in MATLAB or IGOR Pro (WaveMetrics Inc., USA). Statistical analysis was performed in R (www.R-project.org). All the data points across all simulations and all models have been reported and represented in the form of beeswarm, scatter, or box plots in order to avoid any misleading interpretations arising from merely reporting the summary statistics (Marder & Taylor, 2011; Rathour & Narayanan, 2014).

## RESULTS

We built a biophysically and morphologically realistic base model of a CA3b pyramidal neuron, endowed with characteristic active and passive properties (Table 1) adapted from earlier studies (Migliore *et al*., 1999; Lazarewicz *et al*., 2002; Narayanan & Chattarji, 2010) to account for additional constraints on model characteristics (Fig. 1; Table 1). The base model satisfied several signature somato-dendritic characteristics of CA3 pyramidal neurons (Fig. 1; Table 2).

### Multi-parametric multi-objective stochastic search (MPMOSS) yielded a heterogeneous population of CA3 pyramidal neuron models showing characteristic physiology

The use of a single hand-tuned model for analyses and evaluation introduces biases into the overall conclusions. An alternative is to build a population of models, all of which satisfy the characteristic physiological features of the system under consideration (Prinz *et al*., 2003; Marder & Taylor, 2011) and assess the phenomena of interest in the population of models (Fig. 2*A*). Such a population of models, apart from avoiding the obvious biases and disadvantages associated with using a single model for all analyses, also provides a mechanism for assessing heterogeneities and degeneracy in the system under consideration (Prinz *et al*., 2004; Rathour & Narayanan, 2014; Anirudhan & Narayanan, 2015; Srikanth & Narayanan, 2015; Mukunda & Narayanan, 2017; Basak & Narayanan, 2018; Mittal & Narayanan, 2018; Mishra & Narayanan, 2019; Rathour & Narayanan, 2019; Basak & Narayanan, 2020; Jain & Narayanan, 2020; Seenivasan & Narayanan, 2020; Goaillard & Marder, 2021; Roy & Narayanan, 2021; Shridhar *et al*., 2022). To generate a population of models, we employed a multi-parametric multi-objective stochastic search algorithm (Fig. 2*A*) involving 14 parameters (Table 1) and 11 different objectives (Fig. 1; Table 2). We randomly generated 12,000 unique models and found a small subset of 236 models (~1.96%) that satisfied all validation criteria involving characteristic features of CA3 pyramidal neurons (Fig. 2*B–J*; Table 2). These 236 valid models manifested pronounced heterogeneities in their physiological properties, reflective of the heterogeneities observed in CA3 pyramidal neurons (Fig. 2*B–J*; Table 2).

Although we employed 11 different measurements for the validation process, it was essential to confirm that these measurements were indeed assessing different aspects of CA3 pyramidal neuron physiology. Strong correlations between measurements would imply that the validation process did not involve independent measurements but instead validated the same attribute using disparate measurements. To assess this, we computed Pearson’s correlation coefficient among pairs of all physiological measurements across all 236 models (Fig. 3). We found most of these measurements to manifest weak correlations, with only a few pairs showing strong correlations. We found strong correlations between the two sub-threshold excitability measures (*R_in_* and |*Z*|_max_) and between amplitudes of action potentials across the three somato-dendritic locations (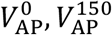, and 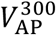). These are expected because the subthreshold excitability measures are dependent on the same parameters, and because the bounds on backpropagating action potentials are with reference to the specific amplitude ranges (of the AP originating at the soma and propagating to different dendritic locations).

**Figure 3:**
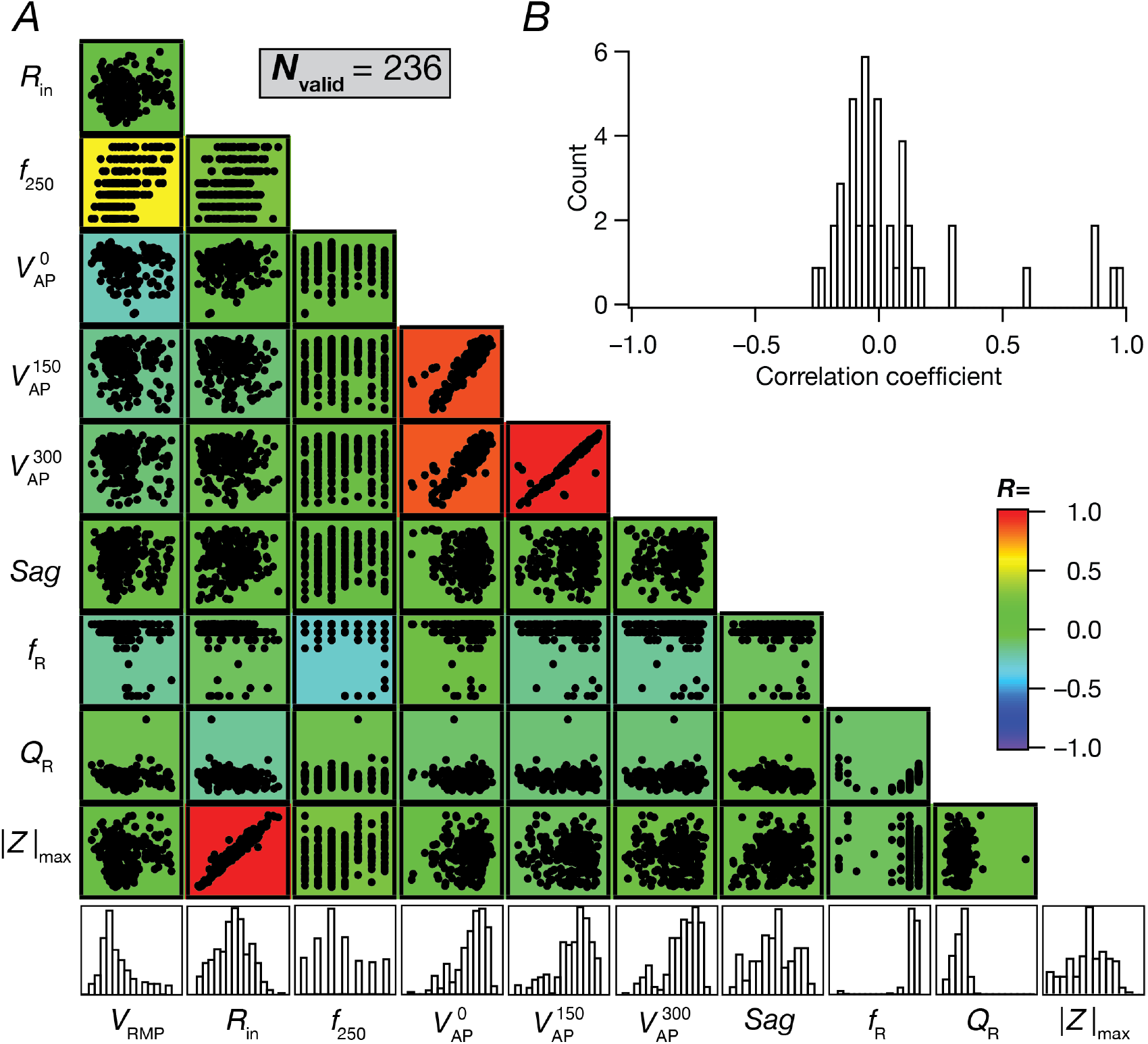
A majority of the pair-wise correlations between the 10 physiological measurements from the 236 valid models were weak. *A*, Scatterplot matrix depicting pairwise relationships between 10 intrinsic measurements from all 236 valid models (*V*_SD_ was omitted from these analyses). Scatter plots are overlaid on a heat map that shows the correlation coefficient for the respective measurement pair. The histograms on the last row indicate the span of each measurement across all valid models. *B*, Histogram of the 45 unique correlation coefficients obtained from the pairwise correlation coefficient matrix shown as a heat map in panel *A*.

### Cellular-scale degeneracy in the manifestation of characteristic physiological properties of CA3 pyramidal neuronal models

Does this CA3 pyramidal neuronal model population require specific parametric values for the emergence of characteristic physiological properties? Could disparate parametric combinations across the parametric space enable the emergence of these signature physiological properties, implying the manifestation of degeneracy? To assess this, we first selected 5 random models out of 236 whose physiologically relevant intrinsic measurements were very similar to each other (Fig. 4*A–F*). Although the physiological measurements were very similar and were characteristic of CA3 pyramidal neurons, the underlying parameters manifested widespread heterogeneities spanning their respective min-to-max ranges (Fig. 4*G*; Table 1). These observations provided us a line of evidence that disparate parametric combinations could come together to elicit similar characteristic physiological properties. The ability of disparate parametric combinations to elicit characteristic physiology has been demonstrated with other neuronal subtypes (Prinz *et al*., 2004; Taylor *et al*., 2009; Rathour & Narayanan, 2014; Rathour *et al*., 2016; Migliore *et al*., 2018; Mittal & Narayanan, 2018; Mishra & Narayanan, 2019; Goaillard & Marder, 2021; Mishra & Narayanan, 2021a). These results show that CA3 pyramidal neurons endowed with signature physiological characteristics could be constructed with disparate parametric combinations.

**Figure 4:**
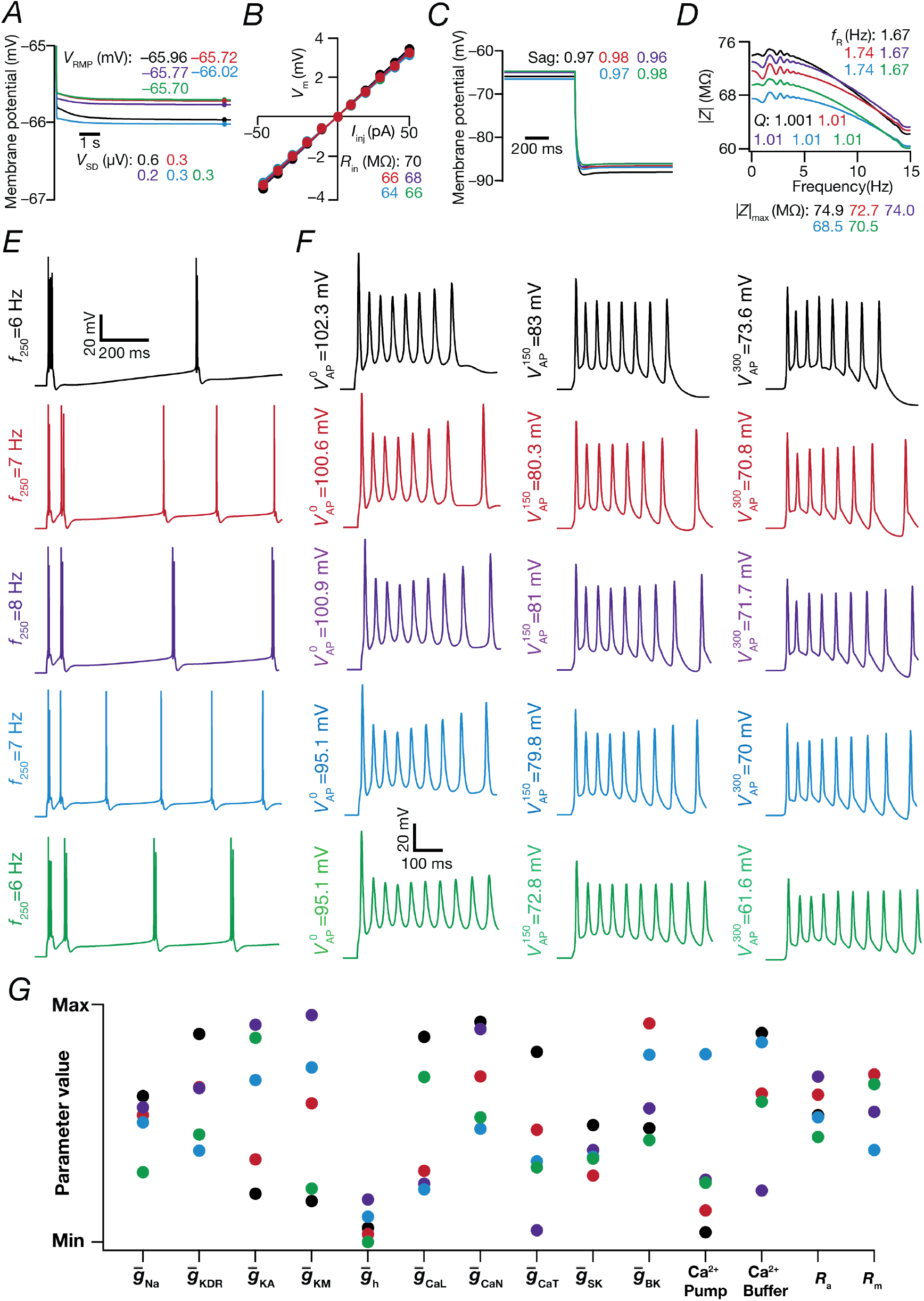
Illustration of degeneracy in the emergence of characteristic physiological measurements in 5 randomly chosen CA3 pyramidal neuron models. *A–F*, Voltage traces and 11 associated intrinsic measurements (Table 2) for 5 distinct valid models chosen from the heterogeneous population of CA3 pyramidal neurons. *G*, Plot representing the normalized parameter values (spanning the respective bounds listed in Table 1) for each of these 5 selected valid models.

These five models provided an illustrative example for the manifestation of cellular scale degeneracy. Were there clusters in the distribution of parameters that governed the 236 valid models? To address this, we plotted the distributions of the 14 parameters that defined these neuronal models and found them to be distributed spanning the range of their respective bounds (Fig. 5*A*). Moreover, these parameters did not manifest strong pairwise correlations across themselves (Fig. 5), suggesting that changes in one parameter were *compensated for* by several different parameters and not just one other parameter. It is important to note that the spread of individual parameters spanning their entire min-to-max range or the lack of strong pairwise correlations among them do not imply that *any* combination of these parameters could yield valid CA3 pyramidal neuron models. It is critical to note that only 236 of the 12,000 models were valid, implying that a large proportion of models (~98.03%) spanning the same parametric range were *rejected* because they did not satisfy the characteristic physiological properties of CA3 pyramidal neurons. Thus, the lack of structure in the underlying parameters should not be considered as evidence that any parametric combination would yield valid CA3 pyramidal neurons. Instead, the lack of structure in the valid parametric space should be interpreted as evidence for the manifestation of degeneracy, whereby disparate (yet specific) *combinations* of parameters yield similar physiological properties (Edelman & Gally, 2001; Prinz *et al*., 2004; Taylor *et al*., 2009; Rathour & Narayanan, 2012, 2014; Anirudhan & Narayanan, 2015; Srikanth & Narayanan, 2015; Rathour *et al*., 2016; Das *et al*., 2017; Basak & Narayanan, 2018; Migliore *et al*., 2018; Mittal & Narayanan, 2018; Mishra & Narayanan, 2019; Rathour & Narayanan, 2019; Basak & Narayanan, 2020; Jain & Narayanan, 2020; Seenivasan & Narayanan, 2020; Goaillard & Marder, 2021; Mishra & Narayanan, 2021a; Roy & Narayanan, 2021; Shridhar *et al*., 2022).

**Figure 5:**
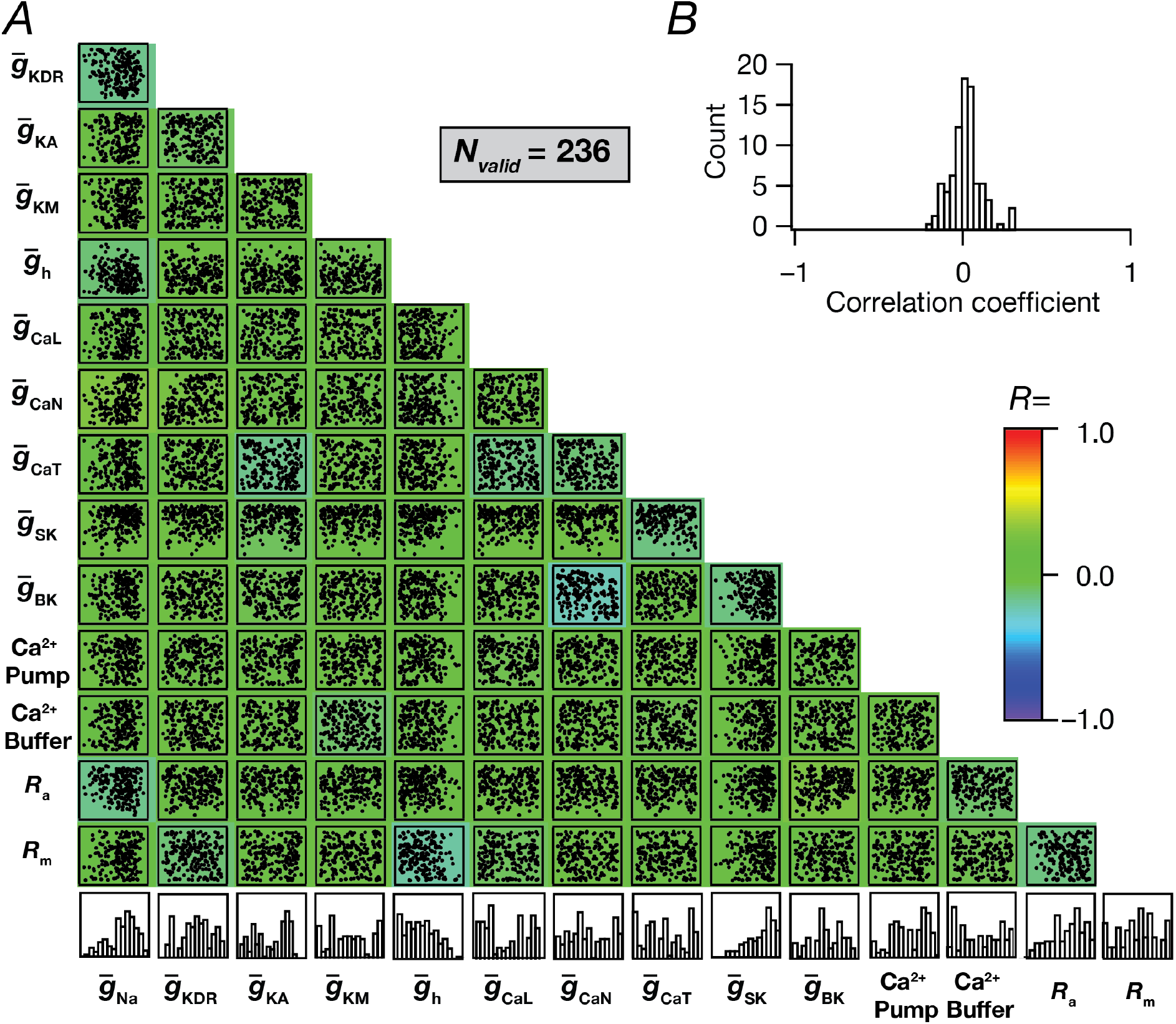
Widespread distributions of individual parameters and weak pairwise correlations between parameters that yielded valid CA3 pyramidal neuron models. *A*, Scatterplot matrix depicting pairwise relationships between all 14 parameters of the 236 valid models. Scatter plots are overlaid on a heat map that shows the correlation coefficient for the respective parameters pair. The histograms on the last row indicate the span of each parameter across all valid models. *B*, Histogram of the 91 unique correlation coefficients obtained from the pairwise correlation coefficient matrix shown as a heat map in panel *A*.

### Heterogeneous CA3 pyramidal neuron population comprised of intrinsically bursting (IB) and regular spiking (RS) models

The CA3 region contains two classes of pyramidal neurons (Wong & Prince, 1978; Hablitz & Johnston, 1981; Traub, 1982; Bilkey & Schwartzkroin, 1990; Traub *et al*., 1991; Migliore *et al*., 1995; Migliore *et al*., 1999; Lazarewicz *et al*., 2002; Hemond *et al*., 2008; Narayanan & Chattarji, 2010; Zeldenrust *et al*., 2018): intrinsically bursting (IB) and regular spiking (RS). IB neurons manifest a burst firing pattern, where action potentials are clustered in distinctly spaced bursts when injected with a depolarizing step current injection. RS neurons elicit action potentials with regular interspike intervals and constant amplitude in response to depolarizing current pulses. If the amplitude of the current stimulus is increased, IB neurons transit to RS firing mode (Migliore *et al*., 1995; Su *et al*., 2001; Narayanan & Chattarji, 2010; Yi *et al*., 2017). We recorded the spiking characteristics of each of the 236 valid models (for a pulse current injection of 240 pA for 5.5 s) and classified them into IB or RS neurons based on their firing profiles and histograms of their interspike intervals, ISI (Fig. 6). Of the 236 valid models, 67 models (~28.38%) were classified as IB neurons (*e.g*., Fig. 6*A–C*; bimodal ISI histograms) and the remaining 169 models (~71.62%) showed regular spiking behavior (*e.g*., Fig. 6*D–F*; unimodal ISI histograms). Consistent with prior observations, we noted that an increase in injected current amplitude in IB neurons resulted in their switch to regular spike firing mode. We performed dimensionality reduction analyses on the active parameters of the 236 models using principal component analysis (PCA) and *t*-distributed stochastic neighbor embedding (*t*-SNE) and visualized the identified RS and IB neurons on the reduced dimensional space (Fig. 7). We found the IB and RS neurons to form clusters with both dimensionality reduction techniques, together suggesting important differences in the parametric space associated with the two classes of neurons (Fig. 7).

**Figure 6:**
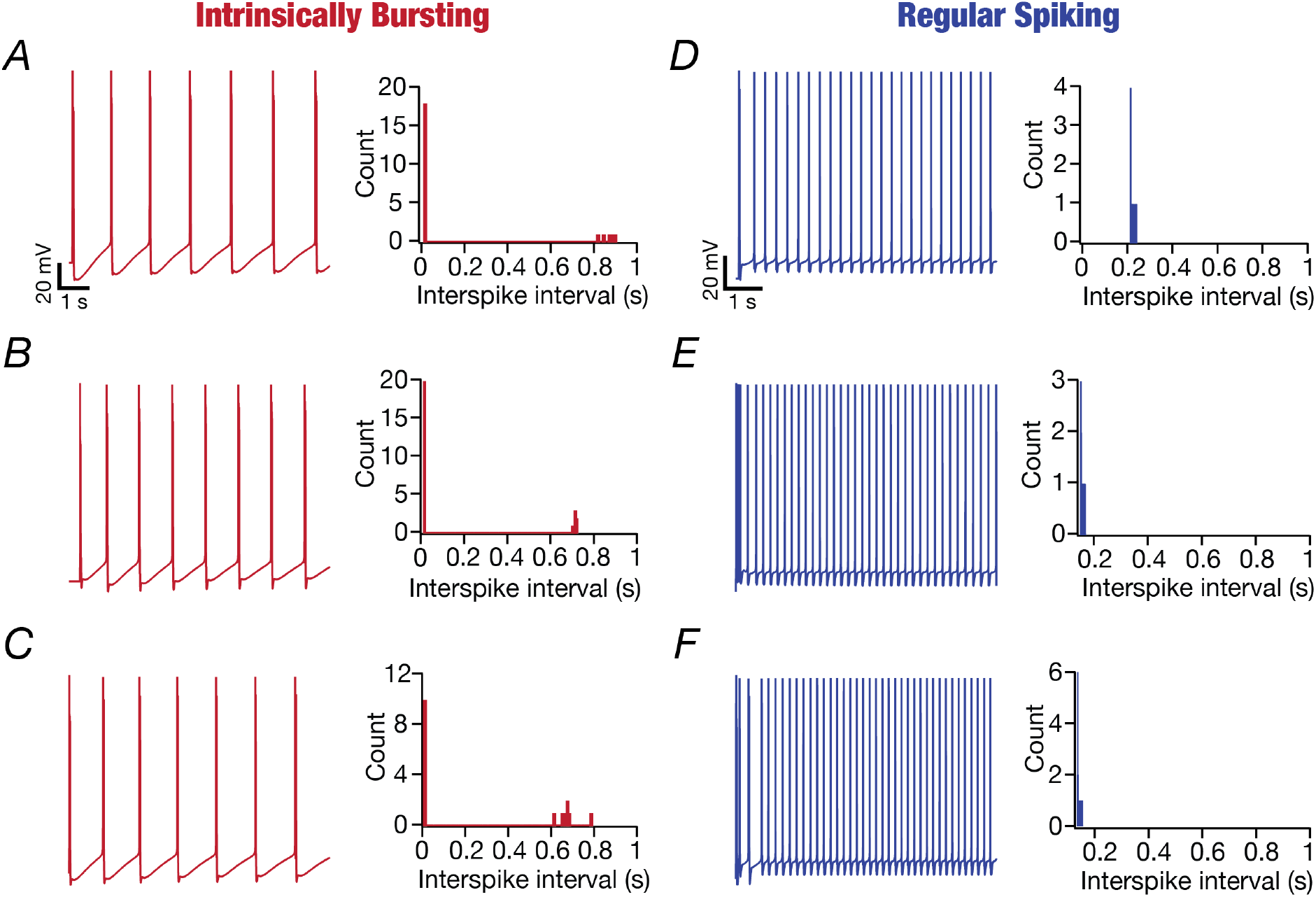
Examples of valid CA3 pyramidal neuron models belonging to the intrinsic bursting and regular spiking classes of firing. All voltage traces in this figure were recorded in response to constant current injection of 240 pA for a duration of 5.5 s. The histograms of the inter-spike intervals (ISI) were derived from the respective traces. *A–C*, Voltage traces from three distinct intrinsically bursting valid models and the associated bimodal ISI histograms. *D–F*, Voltage traces from three distinct regular spiking valid models and the associated unimodal ISI histograms.

**Figure 7:**
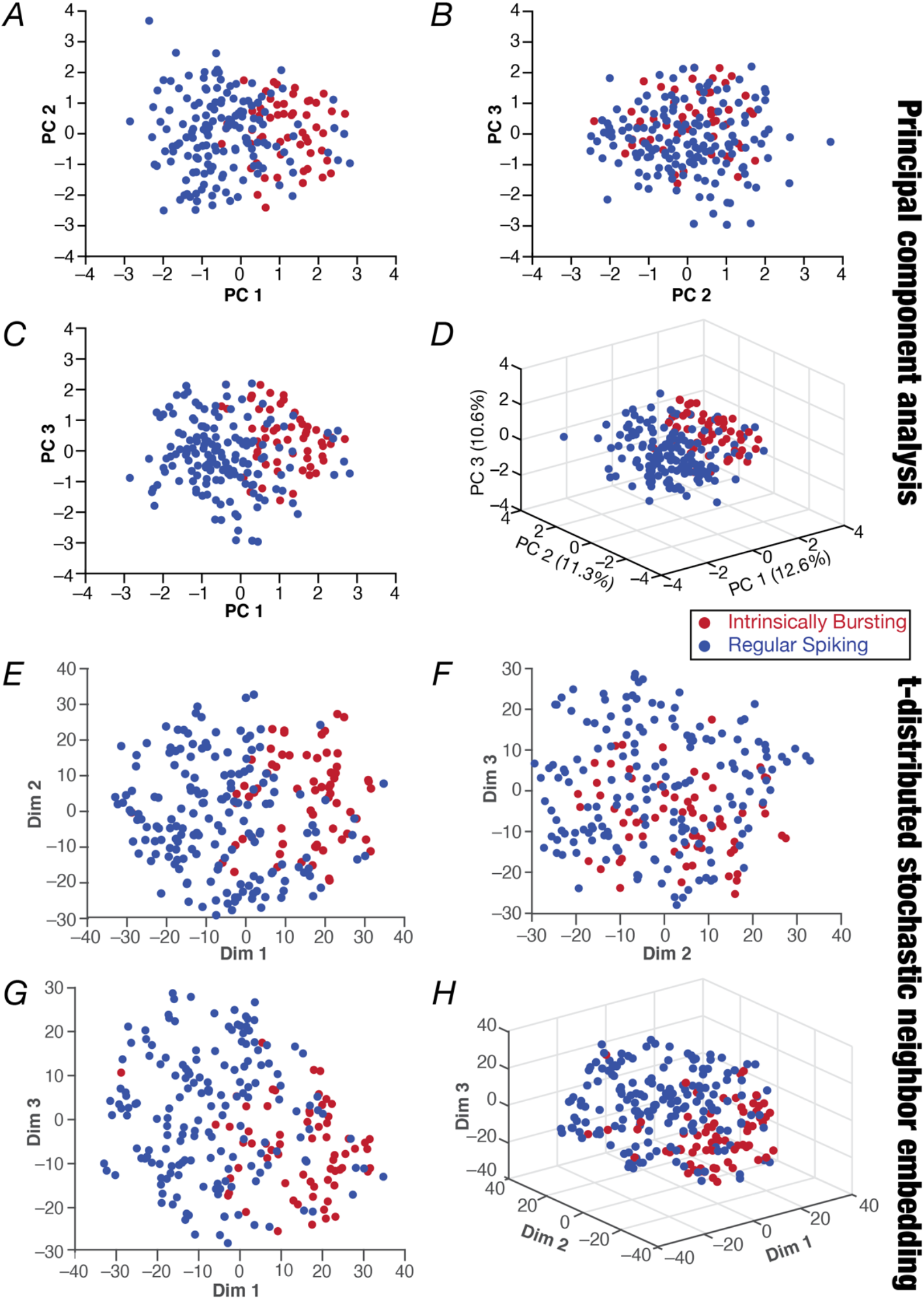
Clustering of regular spiking and intrinsically bursting models in the active parametric space. *A–D*, Principal component analysis performed on the 12 active parameters of the 236 valid models. Shown are pairwise plots between the first three principal dimensions (*A–C*) and a 3D plot showing all three principal components (*D*). The percentage variance explained by each dimension is provided in panel *D* along each axis. *E–H*, Dimensionality reduction results from *t*-distributed stochastic neighbor embedding (*t*-SNE) performed on 12 parameters, showing pairwise (*E–G*) and 3D (*H*) plots.

### Calcium and calcium-activated potassium channels were the dominant contributors to the distinction between intrinsically bursting and regular spiking neuronal classes

Which of the different active parameters contributed to the emergence of two distinct sub-populations of neurons? To address this, we plotted the 5 parameters whose loadings were along the direction of the first principal component (Fig. 8*A*) for all the IB and the RS neurons (Fig. 8*B–F*). Although there were heterogeneities in these parameters across different models, emphasizing the manifestation of degeneracy in the emergence of these two sub-classes, we found significant differences in CaN channel density (Fig. 8*B*) and the density of SK channels (Fig. 8*C*) between IB and RS class of neurons. We observed widespread parametric variability as well as weak pairwise correlations between these 5 parameters in both classes of neurons (Fig. 8*G–H*). In addition, although the density of CaN channels was significantly higher (Fig. 8*B*) and SK channel density was significantly lower (Fig. 8*C*) in IB neurons, we did not find strong negative correlations between these two parameters in IB (Fig. 8*G*) or RS (Fig. 8*H*) neurons. These observations provided further lines of evidence for the critical role of interactions across channel parameters and the manifestation of degeneracy in the emergence of IB or RS class of neurons.

**Figure 8:**
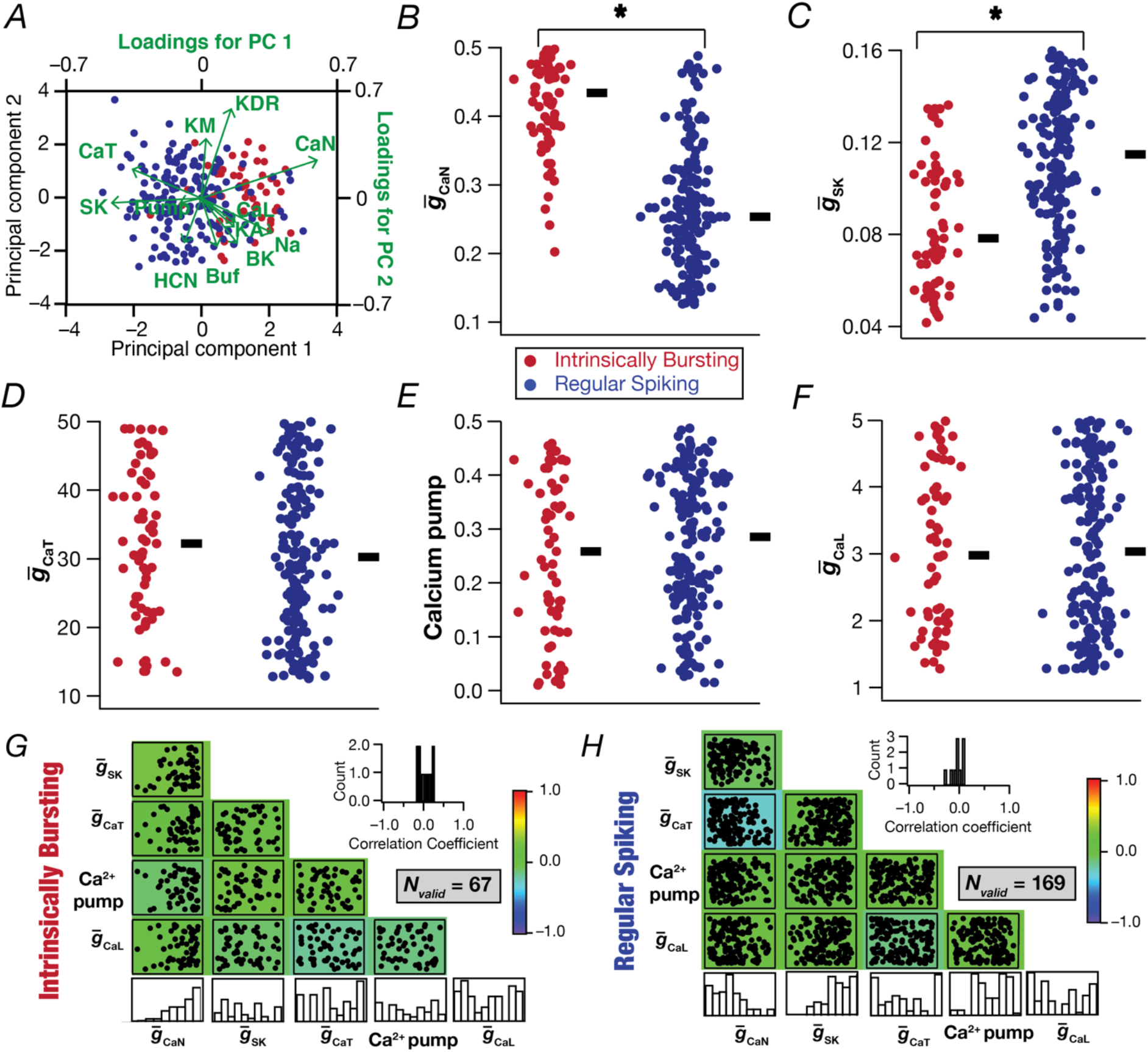
Significant differences in the expression of calcium and calcium-activated potassium channels together contribute to the distinct populations of intrinsically bursting and regular spiking models. *A*, Biplot showing the intrinsically bursting (*N*_valid_ = 67) and regular bursting (*N*_valid_ = 169) models and the projection lines indicating the relationship between the original dimension of the individual variables and their contribution to the relevant principal components. *B–F*, Beeswarm plot showing the distribution of maximal conductance of *N*-type calcium channel (*B*), maximal conductance of calcium-activated potassium channel (*C*), maximal conductance of *T*-type calcium channel (*D*), calcium pump (*E*), and *L*-type calcium channel (*F*) for the intrinsically bursting and the regular spiking populations. Thick lines on the right depict respective median values. *: *p*<0.05, Wilcoxon signed rank test. *G–H*, Scatterplot matrix depicting pairwise relationships between the 5 parameters of the 67 intrinsically bursting (*G*) and the 169 regular spiking (*H*) models. Scatter plots are overlaid on a heat map that shows the correlation coefficient for the respective parameters pair. The histograms on the last row indicate the span of each parameter across the respective set of valid models. The insets represent the histograms of the 10 unique correlation coefficients plotted of the parameter pairs.

### Heterogeneities in complex spike bursting of RS and IB CA3 pyramidal neurons

An important electrophysiological signature of CA3 pyramidal neurons is the manifestation of CSB with various types of inputs arriving along the somato-dendritic arbor (Wong & Prince, 1978; Wong *et al*., 1979; Hablitz & Johnston, 1981; Traub *et al*., 1994a; Lazarewicz *et al*., 2002; Mizuseki *et al*., 2012; Kowalski *et al*., 2016; Oliva *et al*., 2016; Raus Balind *et al*., 2019). We employed the heterogeneous population of 236 model neurons to explore the roles of different ion channels and receptors on the generation of CSB in the CA3b pyramidal neurons. We employed 5 different protocols to elicit CSB (Raus Balind *et al*., 2019): 5 somatic (Fig. 9*A*) current pulses of 600 pA (Fig. 9*B*), 900 pA (Fig. 9*C*), or 1200 pA (Fig. 9*D*); 5 dendritic current pulses of 1000 pA (Fig. 9*E–F*); stimulation of the 100 excitatory synapses with colocalized AMPAR-NMDAR in the *stratum radiatum* through 5 Hz pre-synaptic spike trains (Fig. 9*G–H*). We identified the presence of CSB by placing quantitative constraints on the number of action potentials during the period of current injection or stimulation, the difference between the amplitudes of the first and the second action potentials, and the amplitude of the ramp that was generated during the stimulus duration of each pulse (Table 3). As the first pulse (of the five pulses used for each protocol) invariably elicited valid CSB properties (*e.g*., Fig. 9), we excluded the first pulse from the CSB rate calculation and CSB validation process.

**Figure 9:**
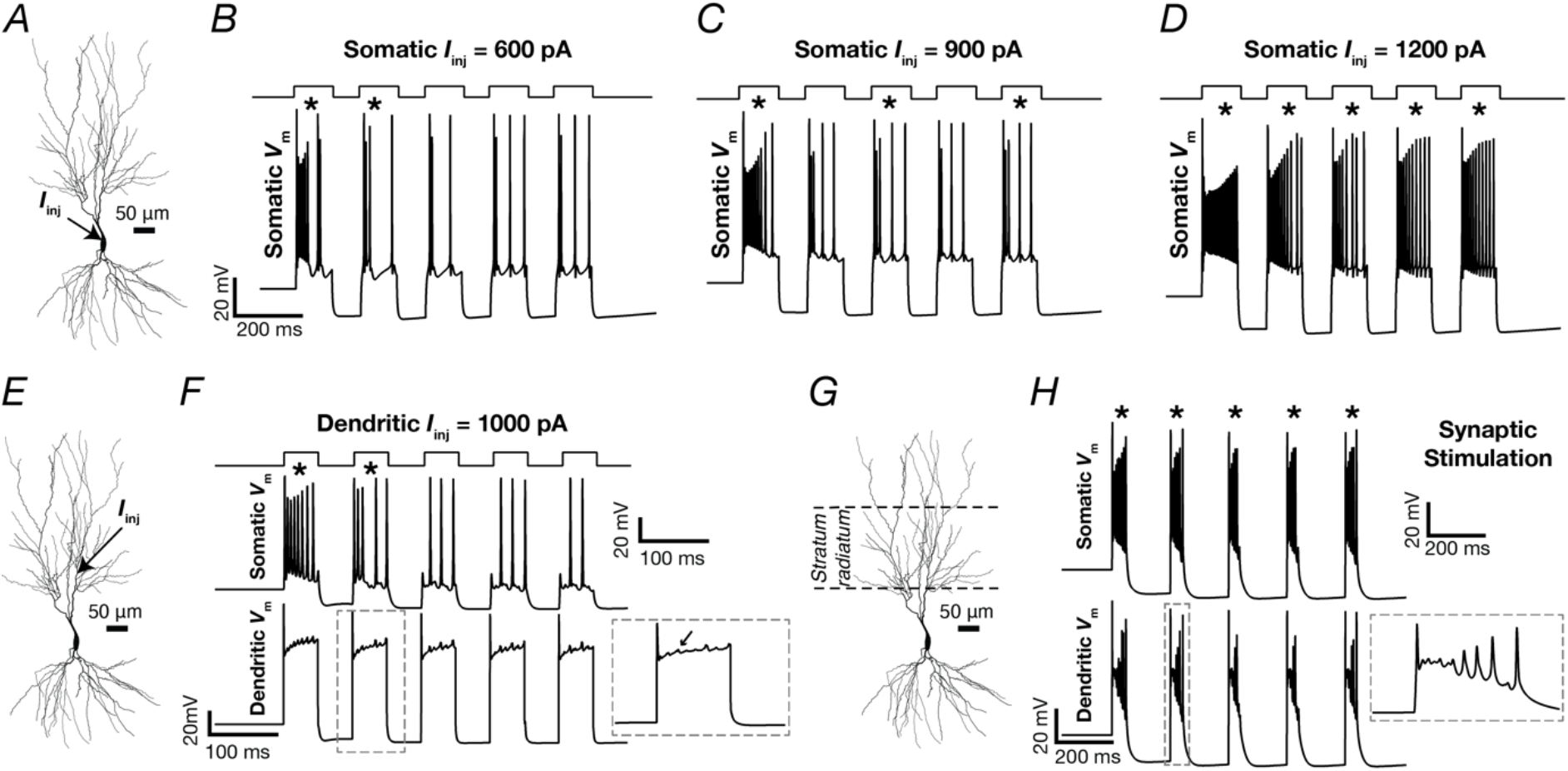
Illustrative examples showing the five different protocols used to assess complex spike bursting in CA3 pyramidal neuron models. *A–D*, A series of 5 current pulses, each of 100 ms duration with inter-pulse interval of 80 ms and amplitude *I*_i*nj*_, was injected into the soma. Depicted are the resultant somatic voltage traces. *I*_i*nj*_=600 pA (*B*), 900 pA (*C*), and 1200 pA (*D*). *E–F*, A series of 5 current pulses, each of 50 ms duration with inter-pulse interval of 50 ms and amplitude *I*_i*nj*_ (1000 pA), was injected into a dendrite located 150 μm away from the soma (*E*). Depicted are the resultant somatic (*F*, *Top*) and dendritic (*F*, *Bottom*, at the location of current injection) voltage traces. Inset (*F*) shows an enlarged view of the dendritic voltage response for the second pulse where a complex spike burst occurred. *H–I*, 100 randomly placed synapses within the *stratum radiatum* (marked in *G*) were stimulated at 5 Hz. Depicted are the resultant somatic (*H*, *Top*) and dendritic (*H*, *Bottom*) voltage traces. Inset (*H*) shows an enlarged view of the dendritic voltage response for the second pulse where a complex spike burst occurred. Asterisks indicate the detection of a valid CSB event satisfying the CSB validity criteria (Table 3).

We repeated the same set of protocols on all 236 valid CA3 pyramidal neuron models and validated models for their ability to manifest CSB using our CSB validation criteria (Table 3). We found pronounced heterogeneity across models in their ability to manifest CSB, as well as in the measurements that were used in their identification (Fig. 10*A–B*). The rate of CSB also showed considerable heterogeneity across models, in a manner that was also dependent on the specific protocol employed (Fig. 10*C*). Overall, CSB propensity increased with increasing somatic current injection (Fig. 10*C*). Whereas the somatic current injection and the synaptic stimulation protocols were efficacious in generating CSB, the dendritic current injection protocol did not produce as many CSB owing to lower ramp voltages and lower differences between the first and the second action potentials (Fig. 10*A–C*). In addition, even within the individual subpopulations of IB and RS models, the CSB measurements manifested considerable and comparable heterogeneities (Fig. 10*D–F*). There were larger proportions of IB (of the 67 valid models) and RS (of the 169 valid models) classes of neurons that showed valid CSB with increasing somatic current intensity, typically with a larger CSB rate as well (Fig. 10*D–F*). Although the 1000 pA dendritic current pulse wasn’t as effective as the somatic current pulses in eliciting CSB, the synaptic stimulation protocol was extremely effective in producing highly reliable CSB (Fig. 10*F*).

**Figure 10:**
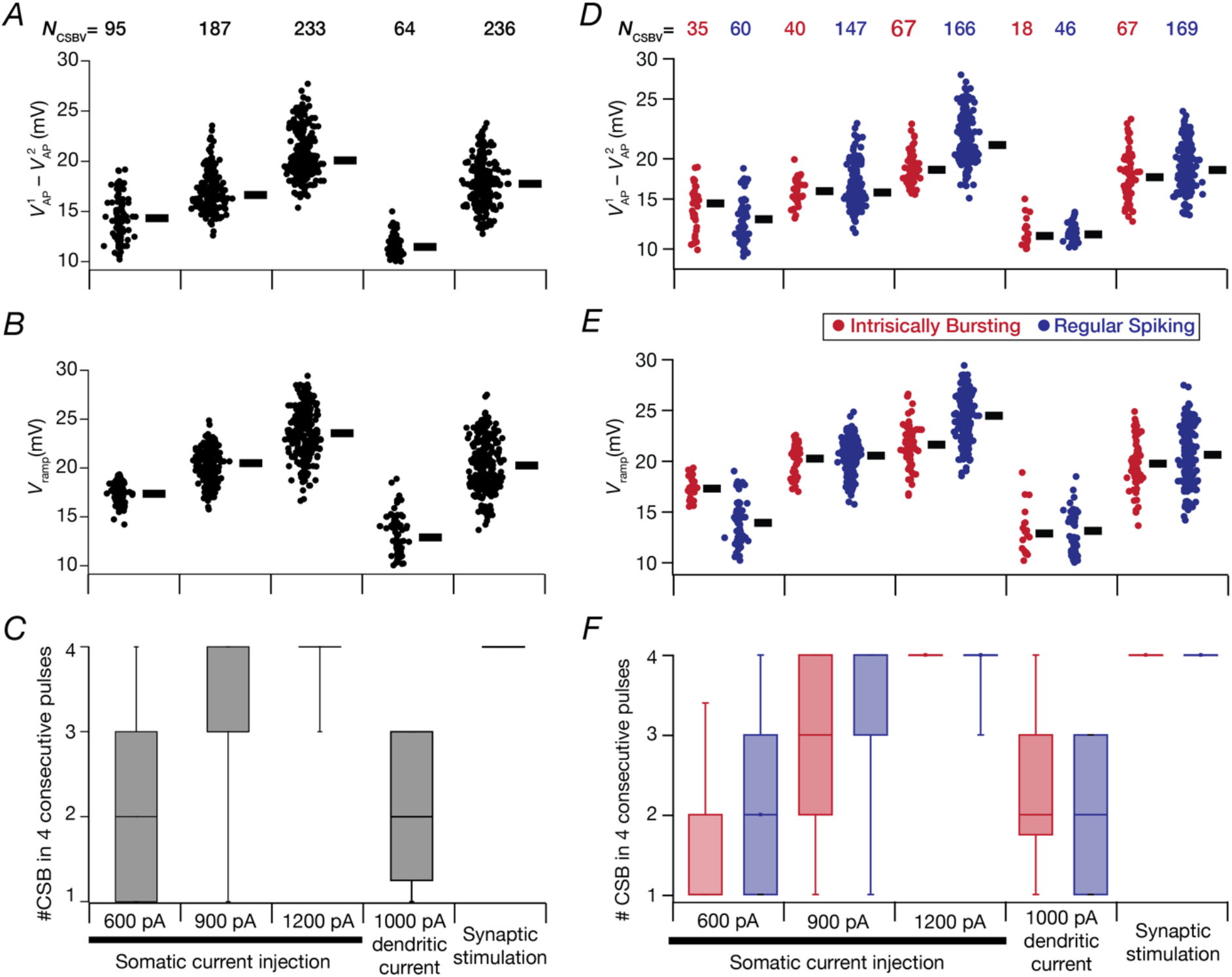
Heterogeneity in CSB measurements across regular spiking and intrinsically bursting models with the five distinct protocols. *A–C*, Distributions of three measurements related to CSB in all models that showed valid CSB: the average difference between the amplitude of the first 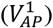 and the second 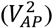 action potentials within a CSB (*A*), average ramp amplitude, *V*_ramp_ (*B*), and the number of CSB (*C*) observed within the 5-pulse protocol (omitting the first of the five pulses). The number of models showing valid CSB (*N*_CSBV_) are provided on the top of panel *A* and hold for panels *A–C*. *D–F*, Same as panels *A–C*, with regular spiking and intrinsically bursting models shown separately.

### Virtual knockout models unveiled synergistic interactions between different channels and receptors in triggering complex spike bursting in CA3 pyramidal neurons

We took advantage of the heterogeneous population of CA3 pyramidal neuron models, manifesting heterogeneous CSB to address the question on the specific role of individual ion channels and receptors in eliciting CSB. To do this, we employed the virtual knockout models (VKM) approach (Rathour & Narayanan, 2014; Anirudhan & Narayanan, 2015; Mukunda & Narayanan, 2017; Basak & Narayanan, 2018; Mittal & Narayanan, 2018; Seenivasan & Narayanan, 2020; Mishra & Narayanan, 2021b; Roy & Narayanan, 2021) by repeating CSB protocols on each of the 236 models in the absence of individual ion channels or receptors. Specifically, we individually set the conductances of the 8 active ion channels (the NaF and KDR channels were not subjected to VKM analyses to allow AP generation across all VKM models) or NMDAR permeability (Nunez *et al*., 1990; Su *et al*., 2001; Sipila *et al*., 2006; Xu & Clancy, 2008; Kim *et al*., 2012; Grienberger *et al*., 2014; Raus Balind *et al*., 2019) to zero in each valid model that showed valid CSB (Table 3) for the two protocols. We repeated the 900 pA somatic current injection (*N_CSBV_*=187 from Fig. 10*A*, implying 187 × 8 = 1496 VKMs for 8 channels) and the 5 Hz synaptic stimulation (*N_CSBV_*=236 from Fig. 10*A*, implying 236 × (8 + 1) = 2124 VKMs for 8 channels and NMDAR) protocols for eliciting CSB in these models. We compared CSB measurements in these VKM models with their respective base models (where all ion channels and receptors were intact) and plotted the percentage changes across all VKMs for each ion channel/receptor (Fig. 11*A–F*).

**Figure 11:**
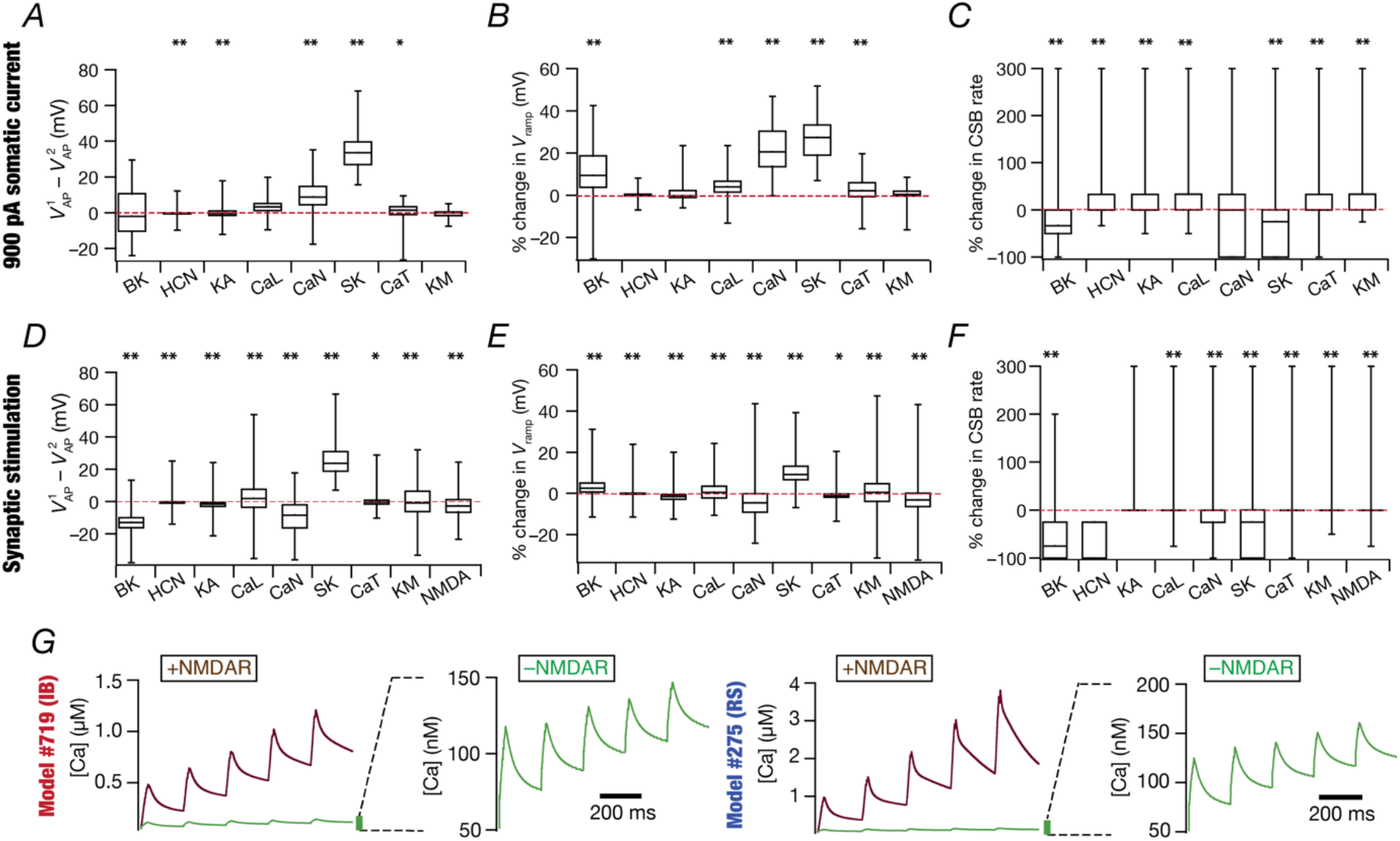
Virtual knockout models unveiled the dominance of calcium, calcium-activated potassium and NMDARs in the emergence of CSB with different protocols. *A–C*, Impact of virtual knockout of different ion channels on all the 187 CSB valid models obtained with 900 pA somatic current injection. Depicted are percentage change induced by the knockout on the average difference between the amplitude of the first 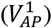 and the second 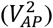 action potentials within a CSB (*A*), average ramp amplitude, *V*_ramp_ (*B*), and CSB rate (*C*). *D–F*, Same as *A–C*, with CSB elicited with synaptic stimulation on 236 models that showed valid CSB. For *A–F*, \**p*<0.05, \*\**p*<0.01, Wilcoxon signed rank test assessed for changes with respect to a ‘no-change’ scenario. *G*, Calcium traces across 5 pulses resulting from stimulation of the *stratum radiatum* synapses for 2 representative models that belong to the IB class (*left*) and RS class (*right*). The impact of virtually knocking out NMDAR (-NMDAR) on calcium is shown, providing a comparison of the calcium traces in the presence of NMDAR (+NMDAR).

We found pronounced heterogeneity in how the deletion of individual channels altered CSB properties (Fig. 11*A–F*). First, eliminating individual ion channels or receptors elicited a large change in certain models, while in other models, the elimination of the same ion channel or receptor had negligible impact (Fig. 11*A–F*). Second, eliminating different ion channels had distinct impacts on the same model, with considerable heterogeneity in the impact of each ion channel. Third, the calcium (CaN) and the calcium-activated potassium (SK) channels had a dominant impact in altering CSB measurements related to both intrinsically (Fig. 11*A–C*) as well as synaptically (Fig. 11*D–F*) elicited CSB. Finally, removal of NMDARs expectedly reduced calcium concentrations at synaptic locations during synaptically-induced CSB, irrespective of whether the models were RS or IB (Fig. 11*G*). Together, these results further emphasized ion-channel degeneracy in the emergence of CSB in CA3 pyramidal neurons, pointing to the role of synergistic interactions between several channels and receptors in CSB emergence.

## DISCUSSION

The principal conclusion of this study is the expression of degeneracy in the emergence of characteristic somato-dendritic physiological properties of CA3 pyramidal neurons, including CSB emergence with different protocols. This degeneracy spanned passive properties, ion-channel expression profiles, as well as calcium-handling mechanisms and from the functional perspective covered several sub- and supra-threshold measurements of soma and dendrites of CA3 pyramidal neurons. These findings therefore eliminate the requirement that a single set of ion channels are essential for CSB generation. Instead, our analyses show that there are several possible combinations of intrinsic mechanisms to elicit CSB, even within the same neuronal subtype, thus providing neurons with multiple degrees of freedom towards CSB generation. These conclusions were arrived at based on a heterogeneous population of CA3 pyramidal neuronal models that were morphologically and biophysically realistic and were validated against several signature electrophysiological properties. Consistent with prior electrophysiological observations, we found two classes of neurons: regular spiking and intrinsic bursting, with calcium and calcium-activated potassium playing critical roles in mediating distinctive firing patterns. We used this heterogeneous population of valid CA3 pyramidal neurons to assess CSB using five distinct protocols and assessed the role of different ion channels and receptors in regulating CSB. Although there was a dominance of calcium and calcium-activated potassium channels in the emergence of CSB, individual deletion of none of the several channels or receptor resulted in the complete elimination of CSB across all models. Together, these analyses provided lines of evidence for the expression of degeneracy in the emergence of CSB, a signature electrophysical characteristic of CA3 pyramidal neurons with several functional roles.

### Mechanisms behind the generation of complex spike bursting

A plethora of mechanisms have been implicated in complex burst generation across different neurons (Wong & Prince, 1978; Hablitz & Johnston, 1981; Migliore *et al*., 1995; Larkum *et al*., 1999; Williams & Stuart, 1999; Wang *et al*., 2000; Bastian & Nguyenkim, 2001; Stuart & Hausser, 2001; Su *et al*., 2001; Lazarewicz *et al*., 2002; Perez-Reyes, 2003; Swensen & Bean, 2003; Krahe & Gabbiani, 2004; Yue & Yaari, 2004; Swensen & Bean, 2005; Sipila *et al*., 2006; Vervaeke *et al*., 2006; Izhikevich, 2007; Cueni *et al*., 2008; Hemond *et al*., 2008; Tazerart *et al*., 2008; Xu & Clancy, 2008; Narayanan & Chattarji, 2010; van Elburg & van Ooyen, 2010; Xu *et al*., 2012; Kadala *et al*., 2015; Morquette *et al*., 2015; Ashhad & Narayanan, 2016; Metzen *et al*., 2016; Condamine *et al*., 2018; Ashhad & Narayanan, 2019). Given the widespread expression of these different ion channels, it is important to recognize that these disparate mechanisms need not be mutually exclusive across different neuronal subtypes. Ion channels with overlapping activation profiles and kinetics can perform the same function of generating these complex bursts. It is important to view neural function from a complex systems perspective, where there are different *functionally segregated* sub-systems (say, ion channels and receptors) that are expressed. These functionally segregated sub-systems then *functionally integrate* to elicit a specific function (say, complex spike bursting). A critical attribute of such a complex system is the ability of multiple combinations of such functionally segregated components to elicit the same function or produce the same output, a phenomenon that has been referred to as degeneracy. Our analyses point to the expression of ion-channel degeneracy in the emergence of CSB in hippocampal neurons, with different ion channels and receptors capable of coming together to contribute to their emergence.

These observations, along with gradients in intrinsic properties along the different anatomical axes imply that one-to-one mapping between individual ion channels or receptors and complex phenomena like CSB generation should be carefully avoided. While it is possible that a specific ion channel has a dominant role in the emergence of CSB in a subset of CA3 pyramidal neurons, a generalization spanning all neurons even within the same anatomical coordinates should be avoided. For instance, our focus here was predominantly on the CA3b neurons, where the density of HCN channels and consequently the manifestation of sag is relatively weak compared to CA3c pyramidal neurons (Raus Balind *et al*., 2019). Therefore, if the validation process involved a large sag and matched with the CA3c pyramidal neurons, the conclusions about the impact of HCN channels on CSB and other intrinsic properties would be different from our conclusions here.

In addition to differences in intrinsic properties, there could be important differences in excitatory and inhibitory inputs and their strengths across the different anatomical axes. It is therefore possible that the CSB propensity could be similar across different neurons because of the balance between the excitatory inputs, the inhibitory inputs and the intrinsic excitability (IE) of the neurons (referred to as E–I–IE balance (Seenivasan & Narayanan, 2020)). Thus, it is extremely important to assess the role of CSB in any given CA3 pyramidal neuron from a holistic perspective that accounts for the morphology of the neuron, the specific intrinsic composition of the neuron (ion channels, pumps, buffer, etc.), the synaptic inputs (excitatory and inhibitory) and their spatial and temporal activation profiles under ethological conditions, and behavioral state dependence involving neuromodulation. The behavioral state-dependence is an important attribute because state-dependent phenomenon like neuromodulation can alter the intrinsic properties and the synaptic inputs onto a neuron thereby providing neurons with context-dependent flexibility of altering their ability to produce CSB (Goldman *et al*., 2001; Tropp Sneider *et al*., 2006; Prince *et al*., 2016; Wagatsuma *et al*., 2018; Alonso & Marder, 2019; Humphries *et al*., 2022).

From a general standpoint, the emergence of a complex spike bursting is *mediated* by interactions between: (i) spike generating conductances (typically NaF and KDR); (ii) a reverberatory positive feedback loop that results in a ramp-like depolarization (mediated by *reverberatory* conductances such as calcium channels, persistent sodium, NMDARs); and (iii) a slower negative feedback loop that ensures that the burst is temporally limited to a few action potential by introducing a hyperpolarization (mediated by slow *restorative* conductances such as SK, KM, HCN). Repetitive bursts could recruit conductances that mediate post inhibitory rebound (such as HCN). In addition to these mediating mechanisms, there are additional *modulating* mechanisms that regulate CSB generation. Such modulating mechanisms could be those governing neural excitability (which regulates CSB by altering the ability of the voltage ramp to elicit action potentials), sources of inhibition that disrupt the voltage ramp or its ability to elicit spikes (such as inhibitory synaptic inputs), and neuromodulation that alters synaptic and/or intrinsic properties of the neuron. Given the several possible combinations of mediating and modulating mechanisms, the ability of *an identical* set of mediating mechanisms could result in CSB in one neuron but not in another depending on the specific modulatory mechanisms expressing in these neurons. In addition, as a neuron could express several of these mediating as well as modulating mechanisms, the ability to manifest similar CSB could arise from disparate combinations of several mechanisms which could vary in a neuron-to-neuron and a context-dependent manner. Thus, it is critical that the global structure of the parametric space spanning all factors that mediate as well as modulate CSB is considered while accounting for heterogeneities as well as degeneracy (Goldman *et al*., 2001; Alonso & Marder, 2019; Rathour & Narayanan, 2019; Goaillard & Marder, 2021). Thus, the generation and analyses of a population of heterogeneous models manifesting degeneracy coupled with experimental recordings that assess CSB under different physiological conditions are essential in studying CSB emergence in an unbiased manner.

### Physiological and pathological implications of degeneracy in the emergence of complex spike bursting

Traditionally, the physiological relevance of bursts has been studied with reference to reliable information transmission, selectivity in transmitted information, and synaptic plasticity (Lisman, 1997; Izhikevich *et al*., 2003; Krahe & Gabbiani, 2004; Metzen *et al*., 2016; Sakmann, 2017). The role of CSB and associated dendritic plateau potentials have received renewed attention with recent demonstrations that have implicated them in behavioral time scale synaptic plasticity (Bittner *et al*., 2015; Bittner *et al*., 2017; Magee & Grienberger, 2020; Zhao *et al*., 2020), perception (Larkum, 2013; Manita *et al*., 2015; Takahashi *et al*., 2016; Takahashi *et al*., 2020), anesthesia (Aru *et al*., 2020; Redinbaugh *et al*., 2020; Suzuki & Larkum, 2020), active sensing (Lavzin *et al*., 2012; Xu *et al*., 2012; Ranganathan *et al*., 2018), and learning (Doron *et al*., 2020; Larkum *et al*., 2022). In addition to these, the significance of bursts under pathological conditions is well established (Traub *et al*., 1994b; Traub *et al*., 1996; McCormick & Contreras, 2001; Su *et al*., 2002; Kole *et al*., 2007; Beck & Yaari, 2008). In this context, our demonstration of degeneracy in the emergence of CSB cautions against a search for a single biophysical mechanism that governs CSB and affects physiological outcomes or rescues pathological conditions, even within neurons of the same subtype. It is critical to account for the heterogeneous expression profiles of different mediating and modulating mechanisms, as well as the ability of disparate combinations of these parameters to elicit similar CSB in executing such a search. The ability of many mechanisms to alter CSB emergence or propensity implies that perturbation to any single mechanism would result in a heterogeneous impact on CSB generation or propensity (Alonso & Marder, 2019; Mishra & Narayanan, 2021b). Thus, in assessing the impact of CSB, their blockade or generation through perturbation of any specific mechanism, it is important to record the heterogeneity across neurons and account for this in the overall analyses. Degeneracy observed in CSB generation, and therefore in its suppression, could also be effectively used for identifying specific drug targets that would be effective in CSB suppression but have minimal secondary and off-target effects.

### Limitations and future directions

In our analyses, we had focused on CA3b pyramidal neurons, constraining models based on recordings from these neurons. However, there are lines of evidence for gradients in intrinsic properties along all anatomical axes: dorsal-ventral, proximal-distal, and deep-superficial (Masukawa *et al*., 1982; Bilkey & Schwartzkroin, 1990; Turner *et al*., 1995; Kowalski *et al*., 2016; Oliva *et al*., 2016; Sun *et al*., 2017; Cembrowski & Spruston, 2019; Raus Balind *et al*., 2019; Sun *et al*., 2020). These differences imply that pyramidal neuron CSB in different parts of the CA3 could be mediated by disparate sets of intrinsic mechanisms, which need careful experimentation and associated populations of heterogeneous computational models across these axes. These analyses should not just account for gradients in biophysical properties, but also account for differences in morphology and afferent synaptic inputs. In addition to these, characterization of ion channel densities as well as gating kinetics along the somato-dendritic axis using cell-attached recordings has been minimal in CA3 pyramidal neurons. It is well established that channel densities and gating kinetics manifest gradients across the somato-dendritic axis of several neuronal subtypes, providing the biophysical basis for their signature electrophysiological properties (Johnston *et al*., 1996; Johnston & Narayanan, 2008; Nusser, 2009; Narayanan & Johnston, 2012; Nusser, 2012). Therefore, it is essential that model populations are refined to incorporate detailed electrophysiological characterization of the different ion channel subtypes in CA3 pyramidal neuron dendrites along each of the anatomical axes mentioned above.

Of the disparate mechanisms that have been implicated in CSB emergence, our study focused on intrinsic properties as well as synaptic activation. However, it is important that the role of different neural circuit mechanisms and heterogeneities therein are assessed towards understanding the emergence of CSB *in vivo*. Prominent among these are pathway interactions between different sets of inputs, the impact of timed inhibitory inputs and E–I balance, as well as astrocytic origins of plateau potentials and bursts. These analyses are best performed in a heterogeneous network of morphologically realistic CA3 pyramidal neuron models, also endowed with interacting populations of heterogeneous interneurons and astrocytes, receiving afferent inputs from the dentate gyrus and the entorhinal cortex. In parallel, detailed electrophysiological experiments associated with each of these different neural-circuit mechanisms involved in CSB emergence should be explored. For instance, while it is known that astrocytic activation results in bursts in other neuronal subtypes, it is important to test this in CA3 pyramidal neurons (Kadala *et al*., 2015; Morquette *et al*., 2015; Ashhad & Narayanan, 2016; Condamine *et al*., 2018; Ashhad & Narayanan, 2019). These heterogeneous network models could then be employed to explore state- and context-dependence using neuromodulatory influence on neural circuit properties as the link. These disparate routes and additional mechanisms in such a model would then provide an ideal substrate for exploring the manifestation of degeneracy, involving all neural circuit components, in the emergence of CSB. Such heterogeneous network models that are explicitly validated against electrophysiological experiments would also provide a substrate for assessing the role of active dendritic properties on LFPs, including the signature sharp wave ripples that are dependent on population bursts in the CA3 region (Sullivan *et al*., 2011; Buzsaki, 2015; Sinha & Narayanan, 2022). Cellular subtypes (neurons and astrocytes) and their intrinsic properties, network connectivity, and synaptic properties are typically very different across different brain regions. Therefore, generalization of conclusions here to CSB in other brain regions should be done after careful exploration of neural-circuit properties and associated heterogeneities in the brain region of interest.

Future analyses could investigate the role of CA3 CSB in learning, plasticity, and behavior using *in vitro* and *in vivo* electrophysiological experiments (Lavzin *et al*., 2012; Xu *et al*., 2012; Larkum, 2013; Bittner *et al*., 2015; Manita *et al*., 2015; Takahashi *et al*., 2016; Bittner *et al*., 2017; Ranganathan *et al*., 2018; Aru *et al*., 2020; Doron *et al*., 2020; Magee & Grienberger, 2020; Redinbaugh *et al*., 2020; Suzuki & Larkum, 2020; Takahashi *et al*., 2020; Zhao *et al*., 2020; Larkum *et al*., 2022) as well as computational analyses involving biophysically constrained plasticity rules dependent on CSB (Milstein *et al*., 2021). As CA3 pyramidal neurons received a multitude of inputs from different brain regions (entorhinal cortex, dentate gyrus, and CA3), it would be interesting to assess CSB emergence through pathway interactions and heterogeneities therein. These experiments, apart from providing further evidence for the importance of CSB in ethological conditions, could also explore routes to harness the manifestation of degeneracy as a tool for designing experiments. Such analyses could explore the role of heterogeneous synaptic distributions, morphological characteristics, and ion-channel distributions on place-cell tuning, dendritic spike generation, and information transmission in CA3 pyramidal neurons through rate and phase codes (Basak & Narayanan, 2018, 2020; Seenivasan & Narayanan, 2020; Roy & Narayanan, 2021). Finally, these experiments could probe the several disparate routes that could regulate CSB generation and propensity towards altering physiological outcomes or reverse pathological conditions.

## ACKNOWLEDGMENTS

The authors thank members of the cellular neurophysiology laboratory for helpful discussions and for comments on a draft of this manuscript. This work was supported by the Wellcome Trust-DBT India Alliance (Senior fellowship to R. N.; IA/S/16/2/502727), the Revati and Satya Nadham Atluri Chair Professorship (R. N.), and the Ministry of education (R. N. & R. R.).

## Author Contributions

R.R. and R.N. designed experiments; R.R. performed experiments; R.R. analyzed data; R.R. and R.N. wrote the paper.

## REFERENCES

Alonso LM & Marder E. (2019). Visualization of currents in neural models with similar behavior and different conductance densities. eLife 8.

Andersen P, Morris R, Amaral D, Bliss T & O’Keefe J. (2006). The hippocampus book. Oxford University Press, New York, USA.

Anirudhan A & Narayanan R. (2015). Analogous synaptic plasticity profiles emerge from disparate channel combinations. J Neurosci 35, 4691–4705.

Aru J, Suzuki M & Larkum ME. (2020). Cellular Mechanisms of Conscious Processing. Trends Cogn Sci 24, 814–825.

Ashhad S & Narayanan R. (2016). Active dendrites regulate the impact of gliotransmission on rat hippocampal pyramidal neurons. Proc Natl Acad Sci U S A 113, E3280–3289.

Ashhad S & Narayanan R. (2019). Stores, Channels, Glue, and Trees: Active Glial and Active Dendritic Physiology. Mol Neurobiol 56, 2278–2299.

Basak R & Narayanan R. (2018). Spatially dispersed synapses yield sharply-tuned place cell responses through dendritic spike initiation. J Physiol 596, 4173–4205.

Basak R & Narayanan R. (2020). Robust emergence of sharply tuned place-cell responses in hippocampal neurons with structural and biophysical heterogeneities. Brain structure & function 225, 567–590.

Bastian J & Nguyenkim J. (2001). Dendritic modulation of burst-like firing in sensory neurons. Journal of neurophysiology 85, 10–22.

Beck H & Yaari Y. (2008). Plasticity of intrinsic neuronal properties in CNS disorders. Nature reviews Neuroscience 9, 357–369.

Bilkey DK & Schwartzkroin PA. (1990). Variation in electrophysiology and morphology of hippocampal CA3 pyramidal cells. Brain Res 514, 77–83.

Bittner KC, Grienberger C, Vaidya SP, Milstein AD, Macklin JJ, Suh J, Tonegawa S & Magee JC. (2015). Conjunctive input processing drives feature selectivity in hippocampal CA1 neurons. Nat Neurosci 18, 1133–1142.

Bittner KC, Milstein AD, Grienberger C, Romani S & Magee JC. (2017). Behavioral time scale synaptic plasticity underlies CA1 place fields. Science 357, 1033–1036.

Buzsaki G. (2015). Hippocampal sharp wave-ripple: A cognitive biomarker for episodic memory and planning. Hippocampus 25, 1073–1188.

Cannon RC, Turner DA, Pyapali GK & Wheal HV. (1998). An on-line archive of reconstructed hippocampal neurons. J Neurosci Methods 84, 49–54.

Cembrowski MS & Spruston N. (2019). Heterogeneity within classical cell types is the rule: lessons from hippocampal pyramidal neurons. Nature reviews Neuroscience 20, 193–204.

Condamine S, Lavoie R, Verdier D & Kolta A. (2018). Functional rhythmogenic domains defined by astrocytic networks in the trigeminal main sensory nucleus. Glia 66, 311–326.

Cueni L, Canepari M, Lujan R, Emmenegger Y, Watanabe M, Bond CT, Franken P, Adelman JP & Luthi A. (2008). T-type Ca2+ channels, SK2 channels and SERCAs gate sleep-related oscillations in thalamic dendrites. Nat Neurosci 11, 683–692.

Das A, Rathour RK & Narayanan R. (2017). Strings on a Violin: Location Dependence of Frequency Tuning in Active Dendrites. Front Cell Neurosci 11, 72.

Doron G, Shin JN, Takahashi N, Druke M, Bocklisch C, Skenderi S, de Mont L, Toumazou M, Ledderose J, Brecht M, Naud R & Larkum ME. (2020). Perirhinal input to neocortical layer 1 controls learning. Science 370.

Edelman GM & Gally JA. (2001). Degeneracy and complexity in biological systems. Proc Natl Acad Sci U S A 98, 13763–13768.

Foster WR, Ungar LH & Schwaber JS. (1993). Significance of conductances in Hodgkin-Huxley models. Journal of neurophysiology 70, 2502–2518.

Frerking M, Schulte J, Wiebe SP & Staubli U. (2005). Spike timing in CA3 pyramidal cells during behavior: implications for synaptic transmission. Journal of neurophysiology 94, 1528–1540.

Goaillard JM & Marder E. (2021). Ion Channel Degeneracy, Variability, and Covariation in Neuron and Circuit Resilience. Annu Rev Neurosci 44, 335–357.

Golding NL, Jung HY, Mickus T & Spruston N. (1999). Dendritic calcium spike initiation and repolarization are controlled by distinct potassium channel subtypes in CA1 pyramidal neurons. J Neurosci 19, 8789–8798.

Goldman DE. (1943). Potential, Impedance, and Rectification in Membranes. The Journal of general physiology 27, 37–60.

Goldman MS, Golowasch J, Marder E & Abbott LF. (2001). Global structure, robustness, and modulation of neuronal models. J Neurosci 21, 5229–5238.

Grienberger C, Chen X & Konnerth A. (2014). NMDA receptor-dependent multidendrite Ca(2+) spikes required for hippocampal burst firing in vivo. Neuron 81, 1274–1281.

Hablitz JJ & Johnston D. (1981). Endogenous nature of spontaneous bursting in hippocampal pyramidal neurons. Cell Mol Neurobiol 1, 325–334.

Harris KD, Hirase H, Leinekugel X, Henze DA & Buzsaki G. (2001). Temporal interaction between single spikes and complex spike bursts in hippocampal pyramidal cells. Neuron 32, 141–149.

Harvey CD, Collman F, Dombeck DA & Tank DW. (2009). Intracellular dynamics of hippocampal place cells during virtual navigation. Nature 461, 941–946.

Hemond P, Epstein D, Boley A, Migliore M, Ascoli GA & Jaffe DB. (2008). Distinct classes of pyramidal cells exhibit mutually exclusive firing patterns in hippocampal area CA3b. Hippocampus 18, 411–424.

Hemond P, Migliore M, Ascoli GA & Jaffe DB. (2009). The membrane response of hippocampal CA3b pyramidal neurons near rest: Heterogeneity of passive properties and the contribution of hyperpolarization-activated currents. Neuroscience 160, 359–370.

Hodgkin AL & Huxley AF. (1952). A quantitative description of membrane current and its application to conduction and excitation in nerve. J Physiol 117, 500–544.

Hodgkin AL & Katz B. (1949). The effect of sodium ions on the electrical activity of giant axon of the squid. J Physiol 108, 37–77.

Holt GR, Softky WR, Koch C & Douglas RJ. (1996). Comparison of discharge variability in vitro and in vivo in cat visual cortex neurons. Journal of neurophysiology 75, 1806–1814.

Humphries R, Mellor JR & O’Donnell C. (2022). Acetylcholine Boosts Dendritic NMDA Spikes in a CA3 Pyramidal Neuron Model. Neuroscience 489, 69–83.

Izhikevich EM. (2007). Dynamical systems in neuroscience: The geometry of excitability and bursting. The MIT Press, Cambridge, MA.

Izhikevich EM, Desai NS, Walcott EC & Hoppensteadt FC. (2003). Bursts as a unit of neural information: selective communication via resonance. Trends Neurosci 26, 161–167.

Jahr CE & Stevens CF. (1990). Voltage dependence of NMDA-activated macroscopic conductances predicted by single-channel kinetics. J Neurosci 10, 3178–3182.

Jain A & Narayanan R. (2020). Degeneracy in the emergence of spike-triggered average of hippocampal pyramidal neurons. Scientific reports 10, 374.

Johnston D, Magee JC, Colbert CM & Cristie BR. (1996). Active properties of neuronal dendrites. Annu Rev Neurosci 19, 165–186.

Johnston D & Narayanan R. (2008). Active dendrites: colorful wings of the mysterious butterflies. Trends Neurosci 31, 309–316.

Kadala A, Verdier D, Morquette P & Kolta A. (2015). Ion Homeostasis in Rhythmogenesis: The Interplay Between Neurons and Astroglia. Physiology 30, 371–388.

Kesner RP. (2007). Behavioral functions of the CA3 subregion of the hippocampus. Learning & memory 14, 771–781.

Kim S, Guzman SJ, Hu H & Jonas P. (2012). Active dendrites support efficient initiation of dendritic spikes in hippocampal CA3 pyramidal neurons. Nat Neurosci 15, 600–606.

Kole MH, Brauer AU & Stuart GJ. (2007). Inherited cortical HCN1 channel loss amplifies dendritic calcium electrogenesis and burst firing in a rat absence epilepsy model. J Physiol 578, 507–525.

Kowalski J, Gan J, Jonas P & Pernia-Andrade AJ. (2016). Intrinsic membrane properties determine hippocampal differential firing pattern in vivo in anesthetized rats. Hippocampus 26, 668–682.

Krahe R & Gabbiani F. (2004). Burst firing in sensory systems. Nature reviews Neuroscience 5, 13–23.

Krichmar JL, Nasuto SJ, Scorcioni R, Washington SD & Ascoli GA. (2002). Effects of dendritic morphology on CA3 pyramidal cell electrophysiology: a simulation study. Brain Res 941, 11–28.

Larkum M. (2013). A cellular mechanism for cortical associations: an organizing principle for the cerebral cortex. Trends Neurosci 36, 141–151.

Larkum ME, Wu J, Duverdin SA & Gidon A. (2022). The Guide to Dendritic Spikes of the Mammalian Cortex In Vitro and In Vivo. Neuroscience 489, 15–33.

Larkum ME, Zhu JJ & Sakmann B. (1999). A new cellular mechanism for coupling inputs arriving at different cortical layers. Nature 398, 338–341.

Lavzin M, Rapoport S, Polsky A, Garion L & Schiller J. (2012). Nonlinear dendritic processing determines angular tuning of barrel cortex neurons in vivo. Nature 490, 397–401.

Lazarewicz MT, Migliore M & Ascoli GA. (2002). A new bursting model of CA3 pyramidal cell physiology suggests multiple locations for spike initiation. Biosystems 67, 129–137.

Lisman JE. (1997). Bursts as a unit of neural information: making unreliable synapses reliable. Trends Neurosci 20, 38–43.

Magee JC. (1998). Dendritic hyperpolarization-activated currents modify the integrative properties of hippocampal CA1 pyramidal neurons. J Neurosci 18, 7613–7624.

Magee JC & Grienberger C. (2020). Synaptic Plasticity Forms and Functions. Annu Rev Neurosci 43, 95–117.

Magee JC & Johnston D. (1997). A synaptically controlled, associative signal for Hebbian plasticity in hippocampal neurons. Science 275, 209–213.

Mainen ZF & Sejnowski TJ. (1996). Influence of dendritic structure on firing pattern in model neocortical neurons. Nature 382, 363–366.

Manita S, Suzuki T, Homma C, Matsumoto T, Odagawa M, Yamada K, Ota K, Matsubara C, Inutsuka A, Sato M, Ohkura M, Yamanaka A, Yanagawa Y, Nakai J, Hayashi Y, Larkum ME & Murayama M. (2015). A Top-Down Cortical Circuit for Accurate Sensory Perception. Neuron 86, 1304–1316.

Marder E & Taylor AL. (2011). Multiple models to capture the variability in biological neurons and networks. Nat Neurosci 14, 133–138.

Masukawa LM, Benardo LS & Prince DA. (1982). Variations in electrophysiological properties of hippocampal neurons in different subfields. Brain Res 242, 341–344.

McCormick DA & Contreras D. (2001). On the cellular and network bases of epileptic seizures. Annual review of physiology 63, 815–846.

Metzen MG, Krahe R & Chacron MJ. (2016). Burst Firing in the Electrosensory System of Gymnotiform Weakly Electric Fish: Mechanisms and Functional Roles. Frontiers in computational neuroscience 10, 81.

Migliore M, Cook EP, Jaffe DB, Turner DA & Johnston D. (1995). Computer simulations of morphologically reconstructed CA3 hippocampal neurons. Journal of neurophysiology 73, 1157–1168.

Migliore M, Hoffman DA, Magee JC & Johnston D. (1999). Role of an A-type K+ conductance in the back-propagation of action potentials in the dendrites of hippocampal pyramidal neurons. J Comput Neurosci 7, 5–15.

Migliore R, Lupascu CA, Bologna LL, Romani A, Courcol JD, Antonel S, Van Geit WAH, Thomson AM, Mercer A, Lange S, Falck J, Rossert CA, Shi Y, Hagens O, Pezzoli M, Freund TF, Kali S, Muller EB, Schurmann F, Markram H & Migliore M. (2018). The physiological variability of channel density in hippocampal CA1 pyramidal cells and interneurons explored using a unified data-driven modeling workflow. PLoS computational biology 14, e1006423.

Miles RM, Le Duigou C, Simonnet J, Telenczuk M & Fricker D. (2014). Recurrent synapses and circuits in the CA3 region of the hippocampus: an associative network. Frontiers in cellular neuroscience 7, 262.

Milstein AD, Li Y, Bittner KC, Grienberger C, Soltesz I, Magee JC & Romani S. (2021). Bidirectional synaptic plasticity rapidly modifies hippocampal representations. eLife 10.

Mishra P & Narayanan R. (2019). Disparate forms of heterogeneities and interactions among them drive channel decorrelation in the dentate gyrus: Degeneracy and dominance. Hippocampus 29, 378–403.

Mishra P & Narayanan R. (2020). Heterogeneities in intrinsic excitability and frequency-dependent response properties of granule cells across the blades of the rat dentate gyrus. Journal of neurophysiology 123, 755–772.

Mishra P & Narayanan R. (2021a). Ion-channel degeneracy: Multiple ion channels heterogeneously regulate intrinsic physiology of rat hippocampal granule cells. Physiol Rep 9, e14963.

Mishra P & Narayanan R. (2021b). Ion-channel regulation of response decorrelation in a heterogeneous multi-scale model of the dentate gyrus. Curr Res Neurobiol 2, 100007.

Mittal D & Narayanan R. (2018). Degeneracy in the robust expression of spectral selectivity, subthreshold oscillations and intrinsic excitability of entorhinal stellate cells. Journal of neurophysiology 120, 576–600.

Mizuseki K, Royer S, Diba K & Buzsaki G. (2012). Activity dynamics and behavioral correlates of CA3 and CA1 hippocampal pyramidal neurons. Hippocampus 22, 1659–1680.

Morquette P, Verdier D, Kadala A, Fethiere J, Philippe AG, Robitaille R & Kolta A. (2015). An astrocyte-dependent mechanism for neuronal rhythmogenesis. Nat Neurosci 18, 844–854.

Mukunda CL & Narayanan R. (2017). Degeneracy in the regulation of short-term plasticity and synaptic filtering by presynaptic mechanisms. J Physiol 595, 2611–2637.

Nakazawa K, Quirk MC, Chitwood RA, Watanabe M, Yeckel MF, Sun LD, Kato A, Carr CA, Johnston D, Wilson MA & Tonegawa S. (2002). Requirement for hippocampal CA3 NMDA receptors in associative memory recall. Science 297, 211–218.

Narayanan R & Chattarji S. (2010). Computational analysis of the impact of chronic stress on intrinsic and synaptic excitability in the hippocampus. Journal of neurophysiology 103, 3070–3083.

Narayanan R & Johnston D. (2007). Long-term potentiation in rat hippocampal neurons is accompanied by spatially widespread changes in intrinsic oscillatory dynamics and excitability. Neuron 56, 1061–1075.

Narayanan R & Johnston D. (2008). The h channel mediates location dependence and plasticity of intrinsic phase response in rat hippocampal neurons. J Neurosci 28, 5846–5860.

Narayanan R & Johnston D. (2010). The h current is a candidate mechanism for regulating the sliding modification threshold in a BCM-like synaptic learning rule. Journal of neurophysiology 104, 1020–1033.

Narayanan R & Johnston D. (2012). Functional maps within a single neuron. Journal of neurophysiology 108, 2343–2351.

Nunez A, Garcia-Austt E & Buno W. (1990). Slow intrinsic spikes recorded in vivo in rat CA1-CA3 hippocampal pyramidal neurons. Experimental neurology 109, 294–299.

Nusser Z. (2009). Variability in the subcellular distribution of ion channels increases neuronal diversity. Trends Neurosci 32, 267–274.

Nusser Z. (2012). Differential subcellular distribution of ion channels and the diversity of neuronal function. Current opinion in neurobiology 22, 366–371.

Oliva A, Fernandez-Ruiz A, Buzsaki G & Berenyi A. (2016). Spatial coding and physiological properties of hippocampal neurons in the Cornu Ammonis subregions. Hippocampus 26, 1593–1607.

Perez-Reyes E. (2003). Molecular physiology of low-voltage-activated t-type calcium channels. Physiol Rev 83, 117–161.

Prince LY, Bacon TJ, Tigaret CM & Mellor JR. (2016). Neuromodulation of the Feedforward Dentate Gyrus-CA3 Microcircuit. Front Synaptic Neurosci 8, 32.

Prinz AA, Billimoria CP & Marder E. (2003). Alternative to hand-tuning conductance-based models: construction and analysis of databases of model neurons. Journal of neurophysiology 90, 3998–4015.

Prinz AA, Bucher D & Marder E. (2004). Similar network activity from disparate circuit parameters. Nat Neurosci 7, 1345–1352.

Ranganathan GN, Apostolides PF, Harnett MT, Xu NL, Druckmann S & Magee JC. (2018). Active dendritic integration and mixed neocortical network representations during an adaptive sensing behavior. Nat Neurosci 21, 1583–1590.

Rathour RK, Malik R & Narayanan R. (2016). Transient potassium channels augment degeneracy in hippocampal active dendritic spectral tuning. Scientific reports 6, 24678.

Rathour RK & Narayanan R. (2012). Inactivating ion channels augment robustness of subthreshold intrinsic response dynamics to parametric variability in hippocampal model neurons. J Physiol 590, 5629–5652.

Rathour RK & Narayanan R. (2014). Homeostasis of functional maps in active dendrites emerges in the absence of individual channelostasis. Proc Natl Acad Sci U S A 111, E1787–1796.

Rathour RK & Narayanan R. (2019). Degeneracy in hippocampal physiology and plasticity. Hippocampus 29, 980–1022.

Raus Balind S, Mago A, Ahmadi M, Kis N, Varga-Nemeth Z, Lorincz A & Makara JK. (2019). Diverse synaptic and dendritic mechanisms of complex spike burst generation in hippocampal CA3 pyramidal cells. Nature communications 10, 1859.

Redinbaugh MJ, Phillips JM, Kambi NA, Mohanta S, Andryk S, Dooley GL, Afrasiabi M, Raz A & Saalmann YB. (2020). Thalamus Modulates Consciousness via Layer-Specific Control of Cortex. Neuron 106, 66–75 e12.

Roy A & Narayanan R. (2021). Spatial information transfer in hippocampal place cells depends on trial-to-trial variability, symmetry of place-field firing, and biophysical heterogeneities. Neural Netw 142, 636–660.

Royer S, Zemelman BV, Losonczy A, Kim J, Chance F, Magee JC & Buzsaki G. (2012). Control of timing, rate and bursts of hippocampal place cells by dendritic and somatic inhibition. Nat Neurosci 15, 769–775.

Sakmann B. (2017). From single cells and single columns to cortical networks: dendritic excitability, coincidence detection and synaptic transmission in brain slices and brains. Experimental physiology 102, 489–521.

Seenivasan P & Narayanan R. (2020). Efficient phase coding in hippocampal place cells. Physical Review Research 2, 033393.

Selinger JV, Kulagina NV, O’Shaughnessy TJ, Ma W & Pancrazio JJ. (2007). Methods for characterizing interspike intervals and identifying bursts in neuronal activity. J Neurosci Methods 162, 64–71.

Shridhar S, Mishra P & Narayanan R. (2022). Dominant role of adult neurogenesis-induced structural heterogeneities in driving plasticity heterogeneity in dentate gyrus granule cells. Hippocampus.

Sinha M & Narayanan R. (2022). Active Dendrites and Local Field Potentials: Biophysical Mechanisms and Computational Explorations. Neuroscience 489, 111–142.

Sipila ST, Huttu K, Voipio J & Kaila K. (2006). Intrinsic bursting of immature CA3 pyramidal neurons and consequent giant depolarizing potentials are driven by a persistent Na+ current and terminated by a slow Ca2+-activated K+ current. The European journal of neuroscience 23, 2330–2338.

Srikanth S & Narayanan R. (2015). Variability in State-Dependent Plasticity of Intrinsic Properties during Cell-Autonomous Self-Regulation of Calcium Homeostasis in Hippocampal Model Neurons. eNeuro 2, ENEURO.0053-0015.2015.

Stuart GJ & Hausser M. (2001). Dendritic coincidence detection of EPSPs and action potentials. Nat Neurosci 4, 63–71.

Su H, Alroy G, Kirson ED & Yaari Y. (2001). Extracellular calcium modulates persistent sodium current-dependent burst-firing in hippocampal pyramidal neurons. J Neurosci 21, 4173–4182.

Su H, Sochivko D, Becker A, Chen J, Jiang Y, Yaari Y & Beck H. (2002). Upregulation of a T-type Ca2+ channel causes a long-lasting modification of neuronal firing mode after status epilepticus. J Neurosci 22, 3645–3655.

Sullivan D, Csicsvari J, Mizuseki K, Montgomery S, Diba K & Buzsaki G. (2011). Relationships between hippocampal sharp waves, ripples, and fast gamma oscillation: influence of dentate and entorhinal cortical activity. J Neurosci 31, 8605–8616.

Sun Q, Jiang YQ & Lu MC. (2020). Topographic heterogeneity of intrinsic excitability in mouse hippocampal CA3 pyramidal neurons. Journal of neurophysiology 124, 1270–1284.

Sun Q, Sotayo A, Cazzulino AS, Snyder AM, Denny CA & Siegelbaum SA. (2017). Proximodistal Heterogeneity of Hippocampal CA3 Pyramidal Neuron Intrinsic Properties, Connectivity, and Reactivation during Memory Recall. Neuron 95, 656–672 e653.

Suzuki M & Larkum ME. (2020). General Anesthesia Decouples Cortical Pyramidal Neurons. Cell 180, 666–676 e613.

Swensen AM & Bean BP. (2003). Ionic mechanisms of burst firing in dissociated Purkinje neurons. J Neurosci 23, 9650–9663.

Swensen AM & Bean BP. (2005). Robustness of burst firing in dissociated purkinje neurons with acute or long-term reductions in sodium conductance. J Neurosci 25, 3509–3520.

Takahashi H & Magee JC. (2009). Pathway interactions and synaptic plasticity in the dendritic tuft regions of CA1 pyramidal neurons. Neuron 62, 102–111.

Takahashi N, Ebner C, Sigl-Glockner J, Moberg S, Nierwetberg S & Larkum ME. (2020). Active dendritic currents gate descending cortical outputs in perception. Nat Neurosci 23, 1277–1285.

Takahashi N, Oertner TG, Hegemann P & Larkum ME. (2016). Active cortical dendrites modulate perception. Science 354, 1587–1590.

Taylor AL, Goaillard JM & Marder E. (2009). How multiple conductances determine electrophysiological properties in a multicompartment model. J Neurosci 29, 5573–5586.

Tazerart S, Vinay L & Brocard F. (2008). The persistent sodium current generates pacemaker activities in the central pattern generator for locomotion and regulates the locomotor rhythm. J Neurosci 28, 8577–8589.

Traub RD. (1982). Simulation of intrinsic bursting in CA3 hippocampal neurons. Neuroscience 7, 1233–1242.

Traub RD, Borck C, Colling SB & Jefferys JG. (1996). On the structure of ictal events in vitro. Epilepsia 37, 879–891.

Traub RD, Jefferys JG, Miles R, Whittington MA & Toth K. (1994a). A branching dendritic model of a rodent CA3 pyramidal neurone. J Physiol 481 (Pt 1), 79–95.

Traub RD, Jefferys JG & Whittington MA. (1994b). Enhanced NMDA conductance can account for epileptiform activity induced by low Mg2+ in the rat hippocampal slice. J Physiol 478 Pt 3, 379–393.

Traub RD, Wong RK, Miles R & Michelson H. (1991). A model of a CA3 hippocampal pyramidal neuron incorporating voltage-clamp data on intrinsic conductances. Journal of neurophysiology 66, 635–650.

Tropp Sneider J, Chrobak JJ, Quirk MC, Oler JA & Markus EJ. (2006). Differential behavioral state-dependence in the burst properties of CA3 and CA1 neurons. Neuroscience 141, 1665–1677.

Turner DA, Li XG, Pyapali GK, Ylinen A & Buzsaki G. (1995). Morphometric and electrical properties of reconstructed hippocampal CA3 neurons recorded in vivo. J Comp Neurol 356, 580–594.

van Elburg RA & van Ooyen A. (2010). Impact of dendritic size and dendritic topology on burst firing in pyramidal cells. PLoS computational biology 6, e1000781.

Vervaeke K, Gu N, Agdestein C, Hu H & Storm JF. (2006). Kv7/KCNQ/M-channels in rat glutamatergic hippocampal axons and their role in regulation of excitability and transmitter release. J Physiol 576, 235–256.

Vickstrom CR, Liu X, Zhang Y, Mu L, Kelly TJ, Yan X, Hu M-m, Snarrenberg ST & Liu Q-s. (2020). T-type calcium channels contribute to burst firing in a subpopulation of medial habenula neurons. eNeuro 7.

Wagatsuma A, Okuyama T, Sun C, Smith LM, Abe K & Tonegawa S. (2018). Locus coeruleus input to hippocampal CA3 drives single-trial learning of a novel context. Proc Natl Acad Sci U S A 115, E310–E316.

Wang SS, Denk W & Hausser M. (2000). Coincidence detection in single dendritic spines mediated by calcium release. Nat Neurosci 3, 1266–1273.

Williams SR & Stuart GJ. (1999). Mechanisms and consequences of action potential burst firing in rat neocortical pyramidal neurons. J Physiol 521 Pt 2, 467–482.

Wolfart J & Roeper J. (2002). Selective coupling of T-type calcium channels to SK potassium channels prevents intrinsic bursting in dopaminergic midbrain neurons. J Neurosci 22, 3404–3413.

Wong R, Prince D & Basbaum A. (1979). Intradendritic recordings from hippocampal neurons. Proc Natl Acad Sci USA 76, 986–990.

Wong RK & Prince DA. (1978). Participation of calcium spikes during intrinsic burst firing in hippocampal neurons. Brain Res 159, 385–390.

Xu J & Clancy CE. (2008). Ionic mechanisms of endogenous bursting in CA3 hippocampal pyramidal neurons: a model study. PloS one 3, e2056.

Xu NL, Harnett MT, Williams SR, Huber D, O’Connor DH, Svoboda K & Magee JC. (2012). Nonlinear dendritic integration of sensory and motor input during an active sensing task. Nature 492, 247–251.

Yi G, Wang J, Wei X & Deng B. (2017). Action potential initiation in a two-compartment model of pyramidal neuron mediated by dendritic Ca(2+) spike. Scientific reports 7, 45684.

Yue C & Yaari Y. (2004). KCNQ/M channels control spike afterdepolarization and burst generation in hippocampal neurons. J Neurosci 24, 4614–4624.

Zeldenrust F, Wadman WJ & Englitz B. (2018). Neural Coding With Bursts—Current State and Future Perspectives. Frontiers in computational neuroscience 12.

Zhao X, Wang Y, Spruston N & Magee JC. (2020). Membrane potential dynamics underlying context-dependent sensory responses in the hippocampus. Nat Neurosci 23, 881–891.

